# Paratenial thalamus engages in reciprocal and broadcast circuits with the prefrontal cortex

**DOI:** 10.64898/2026.03.27.714842

**Authors:** Nigel Dao, Adam G. Carter

## Abstract

The dorsal anterior midline thalamus (aMT) consists of several closely packed nuclei that are important for motivated and emotional behavior. Previous work on aMT has focused on cells and synapses in the paraventricular thalamus (PVT), and little is known about the adjacent paratenial thalamus (PT). Here we examine neural circuits involving PT using a combination of molecular profiling, anatomical tracing, electrophysiology, and optogenetics. We first find that Protein Kinase C-delta (PKCd) selectively labels thalamocortical (TC) cells concentrated in PT but largely absent from neighboring PVT. We show that TC cells in PT project to the infralimbic region (IL) of the medial prefrontal cortex (mPFC), where they contact and drive L2/3 pyramidal cells. In return, we find that IL mPFC primarily projects to PT over nearby PVT, making connections onto reciprocally connected TC cells. However, these cortical inputs are even stronger onto thalamostriatal (TS) and thalamoamygdala (TA) cells, allowing the mPFC to broadcast to the subcortex. Together, our findings help to parcellate aMT, highlight PT as a distinct thalamic nucleus, establish reciprocal connectivity between PT and IL mPFC, and show cortico-thalamic throughput to the subcortex.

## INTRODUCTION

Interactions between the medial prefrontal cortex (mPFC) and dorsal anterior midline thalamus (aMT) govern diverse behaviors and physiological states, including arousal (Gao *et al*., 2020), reward seeking (Otis *et al*., 2017), fear memory (Do-Monte *et al*., 2015), and social behaviors (Yamamuro *et al*., 2020). Studies in rats (Berendse & Groenewegen, 1991; Vertes & Hoover, 2008), cats (Room *et al*., 1985), and primates (Hsu & Price, 2007) indicate that the aMT consists of several small nuclei, including adjacent paraventricular thalamus (PVT) and paratenial thalamus (PT). Previous work has focused on PVT, which makes strong connections with mPFC, nucleus accumbens (NAc), and amygdala. In contrast, very little is known about PT, including the properties of different cell types and their input and output connections with the rest of the brain.

Due to their small size, it has been challenging to distinguish PT and PVT, with borders drawn using cytoarchitecture (Groenewegen & Berendse, 1994) or peptidergic staining (Levey *et al*., 1987; Kirouac, 2015). Genetic markers like calretinin (Calb2) are expressed in both nuclei (Matyas *et al*., 2018; Viena *et al*., 2021), and while neurotensin (Nts) and neurotrophic receptor tyrosine kinase 1 (ntrk1) are expressed in PVT (Li *et al*., 2022; Shima *et al*., 2023), equivalent markers for PT have not been found. Anatomical tracing studies suggest that PT contains abundant thalamocortical (TC) cells, whose axons terminate in L1 and L3 in mPFC (Vertes & Hoover, 2008). However, because of the close proximity of PT and PVT, it also has been difficult to study the functional properties of these connections to the cortex. Identifying nucleus-specific markers thus could better delineate these nuclei and help define their projection neurons and connections.

The aMT also receives long-range afferents from the cortex and a variety of subcortical brain regions (Chen, 1990; Biro *et al*., 2025; Linley *et al*., 2025). Corticothalamic (CT) cells in the mPFC form strong reciprocal connections with multiple thalamic nuclei that support persistent activity (Schmitt *et al*., 2017). For example, we previously found that CT cells in prelimbic (PL) mPFC branch to more caudal thalamic nuclei, including mediodorsal (MD) and ventromedial (VM) thalamus (Collins *et al*., 2018). Anatomical studies also suggest dense mPFC inputs to aMT in multiple species (Room *et al*., 1985; Vertes, 2004; Hsu & Price, 2007). These inputs are thought to drive activity in PVT, which is in turn important for an assortment of motivated behaviors (Kirouac, 2015; McGinty & Otis, 2020; Iglesias & Flagel, 2021). However, the connections made by distinct CT cell classes onto reciprocally connected neurons in PT remain unexplored.

Neurons in aMT project densely to other regions of the limbic system, including the NAc and the basolateral amygdala (BLA) (Chen, 1990; Gimenez-Amaya *et al*., 1995; Vertes & Hoover, 2008; Dong *et al*., 2017; Ma *et al*., 2021). Subcortical projections from the PVT have diverse functional properties, including roles in feeding (Kessler *et al*., 2021), avoidance (Ma *et al*., 2021), valence processing (Li *et al*., 2022), and fear learning (Penzo *et al*., 2015). These cells are also implicated in a host of maladaptive behaviors and neuropsychiatric disorders, including stress (Kooiker *et al*., 2023), pain (Tang *et al*., 2024), and substance use disorders (Zhu *et al*., 2016; Clark *et al*., 2017; Paniccia *et al*., 2024). However, the properties of subcortically projecting cells in PT, their drive by long-range inputs, and their long-range outputs, have not been examined.

We first examine the molecular profile of aMT and identify Protein Kinase C delta (PKCd) as a specific marker for PT. Using virus-based circuit mapping, slice electrophysiology, and optogenetics, we establish that PKCd-expressing (PKCd+) neurons are thalamocortical (TC) cells, which are concentrated in PT and project to infralimbic (IL) mPFC. We show that PKCd+ PT cells receive a range of cortical and subcortical inputs, including prominent inputs from L6 CT cells in IL mPFC. However, we find these CT inputs are heavily biased to PT over PVT, in contrast to previous reports suggesting strong connections between mPFC and PVT. Lastly, we show that these inputs drive AP firing of TC cells but are even stronger at thalamostriatal (TS) and thalamoamygdala (TA) cells. Together, our data reveal novel cells, synapses, and circuits that link the mPFC to the NAc and BLA via the thalamus, identifying PT as a key cortico-thalamo-limbic hub that enables prefrontal cortical control over multiple subcortical networks.

## RESULTS

### PKC delta (PKCd) is a molecular marker for paratenial thalamus

The dorsal anterior midline thalamus (aMT) consists of multiple, closely spaced nuclei, which may have distinct organization and function (Vertes *et al*., 2015). To distinguish between these adjacent areas, we first explored the distributions of known genetic markers, including those previously used to identify different cell types in paraventricular thalamus (PVT). We first used fluorescent *in situ* hybridization (FISH) to label neurons in thin coronal slices (10 µm) for specific genes (**Fig. 1A**). To identify different nuclei, we registered to the Common Coordinate Framework (CCF) using the DAPI signal (Wang *et al*., 2020). We found the commonly used marker Calbindin-2 (*calb2* or calretinin) (Hua *et al*., 2018; Matyas *et al*., 2018; Viena *et al*., 2021; Biro *et al*., 2025) was prominent in PVT, but also extended into the neighboring paratenial thalamus (PT) (*calb2* count: PVT = 126.7 ± 27.5 cells, PT = 118.4 ± 22.2 cells, *p* = 0.6, N = 5 mice) (**Fig. 1B)**. The previously characterized markers neurotrophic receptor tyrosine kinase 1 (*ntrk1*) (Shima *et al*., 2023) and neurotensin (*nts*) (Li *et al*., 2022) primarily labeled PVT over PT (*ntrk1* count: PVT = 81.6 ± 5.8 cells, PT = 2.0 ± 0.6 cells, *p* < 0.0001; *nts* count: PVT = 25.3 ± 7.5 cells, PT = 2.0 ± 0.5 cells, *p* = 0.01; N = 5 mice). In contrast, Protein Kinase C delta (*prkcd* or PKCd) was almost exclusively expressed in PT, with minimal labeling of PVT (*prkcd* count: PVT = 3.4 ± 0.3 cells, PT = 34.6 ± 4.5 cells, *p* < 0.001, N = 5 mice). PT/PVT ratio indicated that *prkcd* expression is 10-fold higher in PT than PVT and higher than all other markers (*calb2 =* 0.9, 95% CI = 0.7-1.1; *nts* = 0.09, 95% CI = 0.06-0.16; *ntkr1* = 0.02, 95% CI = 0.006-0.07; *prkcd* = 9.8, 95% CI = 5.8-16.4; *prkcd* vs. *calb2*: *p* < 0.0001, *prkcd* vs. *nts*: *p* < 0.0001, *prkcd* vs. *ntrk1*: *p* < 0.0001) (**Fig. 1C**). We confirmed this labeling with immunohistochemistry, labeling coronal slices for protein expression, again registering to the CCF (**Fig. 1D)**. We again found that Calb2 protein was expressed in both PVT and PT (PVT = 2319 ± 536 cells, PT = 2145 ± 439 cells, *p* = 0.3, N = 4 mice), whereas PKCd protein was largely restricted to PT (PVT = 62.5 ± 12.1 cells, PT = 481.3 ± 20.5 cells, *p* < 0.001, N = 4 mice; PT/PVT ratio: Calb2 = 0.9, 95% CI = 0.7-1.1, PKCd = 8.0, 95% CI = 4.9-13.2, *p* < 0.001) (**Fig. 1E & 1F**). Together, these results indicate that PKCd is a useful marker for distinguishing PT from the nearby PVT and thereby assessing the cells and connections with other brain regions.

**Figure 1:**
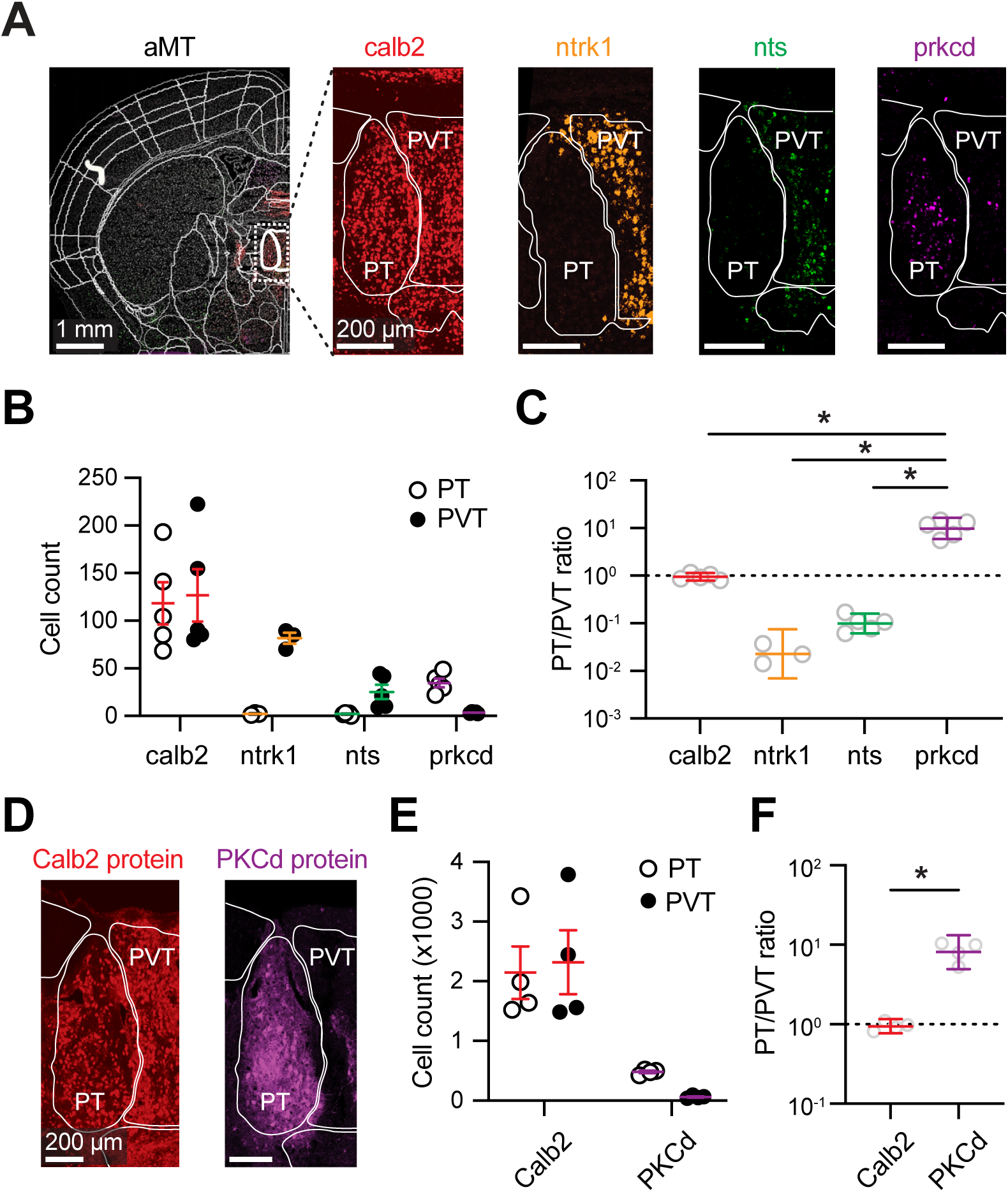
PKCd expression is prominent in paratenial thalamus. **(A)** Confocal images of CCF-aligned coronal slices containing anterior midline thalamus (aMT) (left) and magnification (white dashed line) (right), showing calretinin (*calb2*, red), neutrophin receptor tyrosine kinase 1 (*ntrk1*, orange), neurotensin (*nts*, green) and protein kinase c delta (*prkcd*, magenta) mRNA in paratenial thalamus (PT) and paraventricular thalamus (PVT). Note that ntrk1 comes from a different slice. Scale bars = 1 mm (left) and 200 µm (right). **(B)** Summary of *calb2, ntrk1*, *nts,* and *prkcd-*expressing cell counts in PT and PVT (N = 3-5 mice). **(C)** Summary of *calb2, ntrk1*, *nts,* and *prkcd* PT/PVT ratio. **(D)** Confocal images of immunohistochemically labeled Calb2 (red) and PKCd (magenta) proteins in aMT. Scale bars = 200 µm. **(E)** Summary of Calb2+ and PKCd+ cell counts in PT and PVT (N = 4 mice). **(F)** Summary of Calb2+ and PKCd+ PT/PVT ratio. Values are mean ± SEM or geometric means ± 95% CI (panels C & E). * = p<0.05

### Thalamocortical cells express PKCd in paratenial thalamus

Neurons in the dorsal aMT send outputs to several targets, including medial prefrontal cortex (mPFC), nucleus accumbens (NAc), and basolateral amygdala (BLA). We next used retrograde labeling to assess the distributions of these different projection neurons in PT and PVT. We found injections of rabies virus expressing GFP (RV-GFP) into the mPFC labeled presynaptic thalamocortical (TC) cells primarily in PT, with minimal labeling in nearby PVT (PVT = 195.6 ± 32.8 cells, PT = 549.0 ± 73.5, *p* < 0.0001, N = 6 mice) (**Fig. 2A & 2B**). To rule out any tropism for PT or PVT projection neurons, we also performed retrograde tracing using RV, AAV-retro (AAVrg), and cholera toxin subunit B (CTB). Similar preferential labeling of PT was observed with each method, although AAVrg labeled fewer cells in thalamus (**Fig. S1A & S1B**). Staining for PKCd confirmed location within PT and indicated that most TC cells are PKCd positive (PKCd+) (**Fig. 2C & S1C**). In contrast, injections into NAc or BLA labeled thalamostriatal (TS) and thalamoamygdala (TA) cells in both PT and PVT, with more labeling in the latter (TS: PVT = 491.8 ± 38.6 cells, PT = 238.2 ± 35.7 cells, *p* = 0.007, N = 5 mice; TA: PVT = 340.4 ± 43.3 cells, PT = 115.8 ± 28.8 cells, *p* < 0.001, N = 5 mice; PT/PVT ratio: TC = 2.8, 95% CI = 2.2-3.5; TS = 0.4, 95% CI = 0.3-0.7; TA = 0.3, 95% CI = 0.2-0.4, *p* < 0.0001) (**Fig. 2A & 2B**). Neither TS nor TA cells showed any overlap with PKCd labeling, indicating that they are a separate population (PKCd+ co-labeled count: TC = 274.5 ± 32.6 cells, TS = 7.8 ± 2.3 cells, TA = 6.6 ± 1.1 cells, TC vs. TS: *p* < 0.0001, TC vs. TA: *p* < 0.0001) (**Fig. 2C & S1C)**. Dual retrograde injections into NAc and BLA of the same animals showed some co-labeling of cells in both PT and PVT, suggesting at least a fraction of these neurons can bifurcate to both subcortical targets (**Fig. 2D**). All three projection neurons were co-labeled with Calb2 (**Fig. S1D & S1E**), in accordance with previous reports suggesting Calb2+ cells in aMT project broadly to multiple structures (Matyas *et al*., 2018).

**Figure 2:**
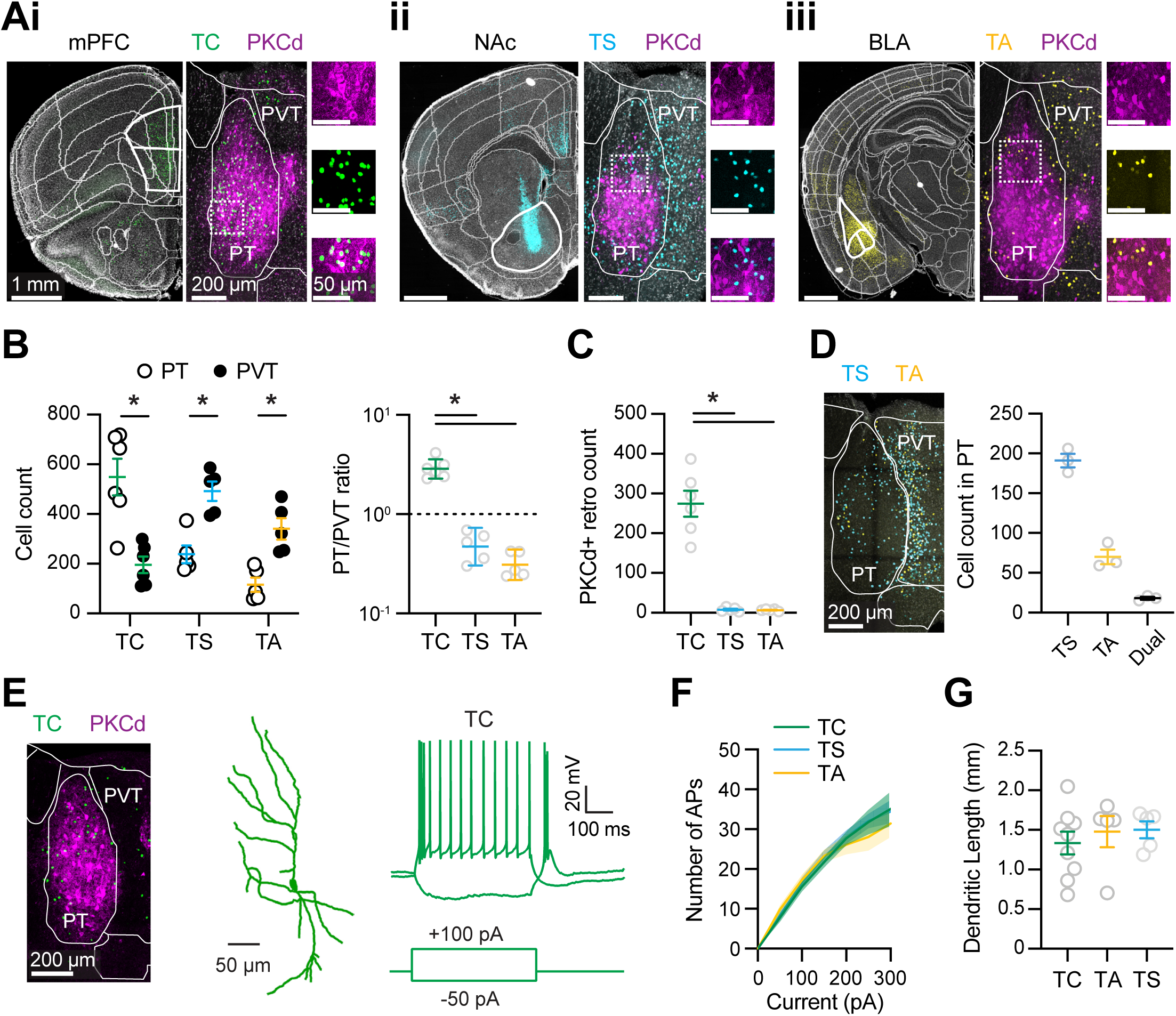
Characterizing projection neurons in paratenial thalamus. **(A)** *Left*, Images of RV-H2B-GFP injections into mPFC (Ai, green), NAc (Aii, cyan), and BLA (Aiii, yellow), with injection sites outlined in white. *Middle*, Magnified confocal images showing retrogradely labeled TC, TS and TA cells in PT and PVT, overlaid with PKCd IHC (purple). *Right*, Further magnified confocal images showing PKCd IHC (top), RV-labeled cells (middle), and merged images (bottom). Scale bars = 1 mm, 200 µm, and 50 µm. **(B)** Summary of retrogradely labeled TC, TS, and TA cell counts in PT and PVT (left) and PT/PVT ratio (right) (N = 5-6 mice per cell type). **(C)** Summary of TC, TS, and TA cells co-labeled with PKCd in PT (N = 5-6 mice per cell type). **(D)** *Left*, Confocal image of TS (blue) and TA (yellow) cell labeling in aMT. Scale bar = 200 µm. *Right*, Summary of TS-only, TA-only, and TS/TA dual-labeled cells in PT (N = 3 mice). **(E)** Representative image of AAVrg-labeled TC cells (green) in PT identified by PKCd IHC (purple) (left), dendritic reconstruction (middle) and membrane potential responses to current injections (right). Scale bars = 200 µm, 50 µm and 20 mV x 100 ms. **(F)** Summary of frequency-current (F-I) curves (n = 8-9 cells, N = 4-5 mice). **(G)** Summary of total dendritic lengths (n = 5-9 cells, N = 4-6 mice). Values are mean ± SEM or geometric means ± 95% CI (panel B). * = p<0.05 *(see also Figures S1 & S2)*

In separate experiments, we also explored the morphological and physiological properties of TC, TS, and TA cells in PT, as well as TC cells in PVT. After injecting retrograde tracers, we prepared acute brain slices and made whole-cell current-clamp recordings from labeled cells (**Fig. 2E**). We injected current steps to assess AP firing and passive properties and filled cells with Alexa-594 to image morphology. We found all three cell types fired APs in response to positive current injections and displayed rebound firing to negative current injections (**Fig. 2F**), similar to TC cells in other parts of thalamus (Llinás & Jahnsen, 1982). We also found that all three cell types had characteristic radial dendrites (**Fig. 2G**), which were broadly similar to TC cells in other parts of thalamus (McAllister & Wells, 1981; Harris, 1986). We found minimal differences in the resting or passive properties of these cells, but did observe subtle differences in rebound AP firing after hyperpolarization (**Fig. S2A & S2B**). We further quantified dendritic morphology with Sholl analysis and found no differences between these cell types (**Fig. S2C**). These findings indicate that cells in PT have morphological and physiological properties similar to other parts of thalamus.

### PKCd+ cells in paratenial thalamus engage the infralimbic mPFC

To explore the connectivity of PKCd+ cells in PT, we next injected AAV-DIO-YFP into the aMT of PKCd-Cre mice (N = 5 mice) that express Cre under the PKCd promoter (Haubensak *et al*., 2010). YFP-expressing cells overlapped with PKCd labeling in PT but not PVT (**Fig. 3A**), consistent with our FISH and IHC labeling experiments, and confirming the specificity of this mouse line. We looked for green axons in serial coronal sections of the brain, and observed prominent labeling in several frontal cortical areas, with the most labeling in infralimbic cortex (IL) (**Fig. 3B & 3C**). In contrast, we observed minimal axonal labeling in subcortical areas that are potential targets of aMT, including NAc and BLA (**Fig. 3B & 3C**), consistent with the lack of PKCd overlap in TS and TA cells in our retrograde labeling experiments. Quantifying fluorescence across layers of the IL indicated that axons were concentrated in both cortical layers 1 and 2/3 (**Fig. 3D**), similar to equivalent inputs from mediodorsal (MD) thalamus (Collins et al., 2018), but in contrast to inputs from ventromedial (VM) thalamus (Cruikshank *et al*., 2012; Anastasiades *et al*., 2021). Interestingly, we also noticed that labeling of PKCd+ axons closely overlapped with PKCd IHC in the mPFC (**Fig. 3D**), and that PKCd IHC densely labels thalamo-recipient layers in many other parts of cerebral cortex (**Fig. S3A**), suggesting that this may be a broad marker of thalamocortical connections across the brain. Intrinsic properties and dendritic morphology of these PKCd+ cells resemble those of retrogradely labeled TC cells (n = 8 cells, N = 3 mice) (**Fig. S3B-E**). These findings confirm PKCd+ cells in aMT are primarily restricted to PT and indicate they are TC cells.

**Figure 3:**
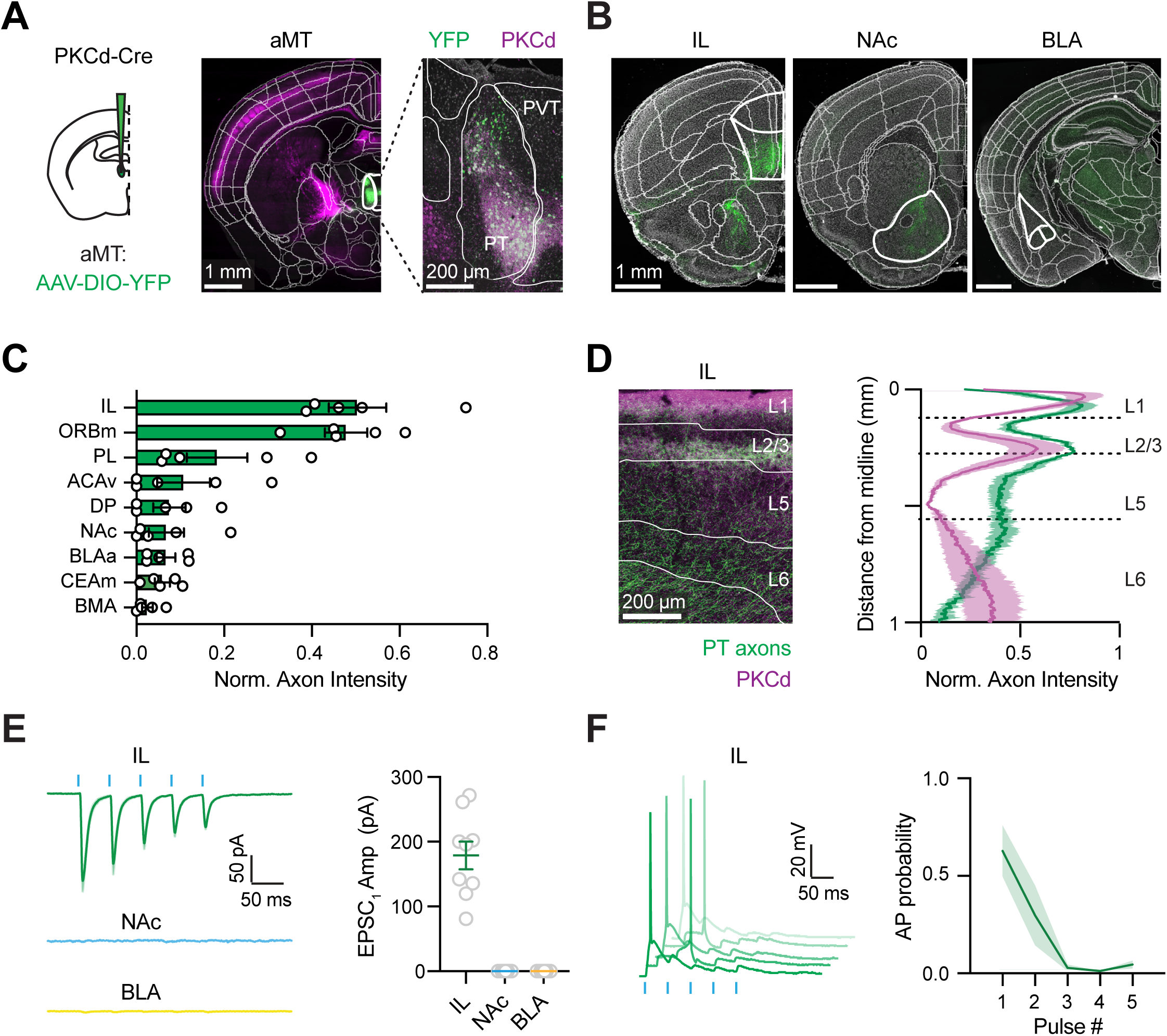
PKCd+ cells in paratenial thalamus drive the infralimbic mPFC. **(A)** *Left*, Schematic of AAV-DIO-YFP injection into the anterior midline thalamus (aMT) of PKCd-Cre mice. *Middle*, CCF-aligned coronal slice showing YFP labeling (green) and PKCd labeling (purple). *Right*, Magnification of PT and PVT, showing labeled cells only in PT. Scale bars = 1 mm and 200 µm. **(B)** Coronal slices showing PKC+ axon labeling in IL (left) and absence of labeling in NAc (middle) and BLA (right). Areas of interest are outlined in white. Scale bars = 1 mm. **(C)** Summary of PKCd+ axon distributions across cortical and subcortical areas (N = 5 mice). **(D)** *Left,* Confocal image of PKC+ axons (green) and PKCd labeling (purple) in IL. *Right,* summary of PKC+ axons and PKCd protein expression in IL (N = 5 mice). **(E)** *Left*, Average PKC+ cell-evoked EPSCs at L2/3 pyramidal neurons in IL (top), MSNs in NAc medial shell (middle), and pyramidal cells in BLA (bottom). Scale bars = 50 pA x 50 ms. *Right,* Summary of EPSC_1_ amplitudes in IL, NAc, and BLA (n = 7-9 cells per area, N = 5 mice). Note that NAc and BLA values are approximately zero and overlap with the y-axis. **(F)** *Left*, PKC+ cell-evoked firing of L2/3 pyramidal cells in IL, showing 5 separate trials. Scale bars = 20 mV x 50 ms. *Right*, Summary of firing rate as a function of stimulation pulse number (n = 6 cells, N = 5 mice). Values are means ± SEM. *(see also Figure S3)*

To explore functional connections from PT to IL and other areas, we next injected AAV-DIO-ChR2 into the aMT of PKCd-Cre mice. After waiting for expression and transport of virus, we prepared acute coronal slices of IL, NAc, and BLA. We used optogenetics to stimulate presynaptic inputs with trains of light, which trigger presynaptic release even from severed axons (5 pulses, 2 ms duration, 10 Hz frequency, 5-10 mW power) (Petreanu *et al*., 2009). We found that repetitive stimulation evoked prominent, depressing EPSCs at L2/3 pyramidal cells in IL (EPSC_1_ = 178.8 ± 21.4 pA, PPR (EPSC_5_/EPSC_1_) = 0.3 ± 0.0, n = 9 cells, N = 6 mice) (**Fig. 3E & S3F**), reminiscent of MD but not VM inputs to the prelimbic cortex (PL) (Collins et al., 2018; Anastasiades et al., 2021; Kamalova et al., 2024). In contrast, PKCd+ PT inputs evoked minimal EPSCs at medium spiny neurons (MSNs) in the NAc (EPSC_1_ = 0.04 ± 0.03 pA, n = 8 cells, N = 6 mice) and pyramidal cells in the BLA (EPSC_1_ = 0.09 ± 0.02 pA, n = 8 cells, N = 6 mice) (**Fig. 3E & S3F**). Additionally, optical stimulation evoked direct excitation at E_GABA_ (-60 mV) and feedforward inhibition at E_Glu_ (+20 mV) with a delay of 6.6 ± 1.2 ms (n = 7 cells) (**Fig. S3G**). In separate current-clamp recordings, repetitive stimulation also evoked strong but transient firing of L2/3 pyramidal cells, similar to MD but not VM inputs (AP probability at pulse 1 = 0.6 ± 0.1; n = 6 cells, N = 5 mice) (**Fig. 3F**). Together, these results indicate that PKCd+ cells in PT are TC cells that contact and drive activity at L2/3 pyramidal cells in IL, but not cells in subcortical areas like NAc and BLA.

### Paratenial thalamus receives robust inputs from infralimbic mPFC

Neurons in aMT are known to receive long-range afferents from several other brain regions, but most studies have focused on PVT (Chen, 1990; Hsu & Price, 2007; Li et al., 2024). To begin to identify long-range inputs specifically onto PT cells, we next used Cre-dependent rabies tracing (Callaway & Luo, 2015). In PKCd-Cre mice, we first injected AAV helper viruses to express the G glycoprotein and the TVA receptor. After waiting 2-3 weeks for expression, we then injected EnvA-pseudotyped RV virus in PT to express histone-tagged GFP in presynaptic cells (**Fig. 4A**). Eight days later, we sectioned the brain, aligned to the CCF, and detected fluorescent starter cells in PT. We also used semi-automated software to detect presynaptic input cells residing in different locations across the brain (n = 57,883 ± 7,515 cells, N = 4 mice) (**Fig. 4B, Fig. S4A & S4B**). We found most presynaptic input cells are concentrated in frontal cortex (28.5 ± 3.4% of total presynaptic cells), with additional but fewer input cells in hypothalamus (13.8 ± 1.7%), septum (9.4 ± 1.2%) and ventral hippocampus (7.1 ± 1.0%) (**Fig. 4C**). We observed no presynaptic input cells in several other subcortical regions that are innervated by aMT, including the NAc and BLA (Dong *et al*., 2017; Ma *et al*., 2021; Tang *et al*., 2024). Within cortex, most presynaptic input cells resided in mPFC (31.5 ± 5.6% of total cortical inputs), including PL and IL, with additional but fewer cells in orbitofrontal cortex (ORBm, 15 ± 1.5%) and anterior insular cortex (aIC, 15.3 ± 2.2%) (**Fig. 4C**). Within IL, we found that most presynaptic input cells were in layer 6 (L6), with a relatively small number of cells in layer 5 (L5) (**Fig. 4D & S4C**). Together, these results indicate that mPFC is a major input to the PT but not the adjacent PVT in dorsal aMT, suggesting the possibility of a reciprocal cortico-thalamo-cortical loop linking these two brain regions.

**Figure 4:**
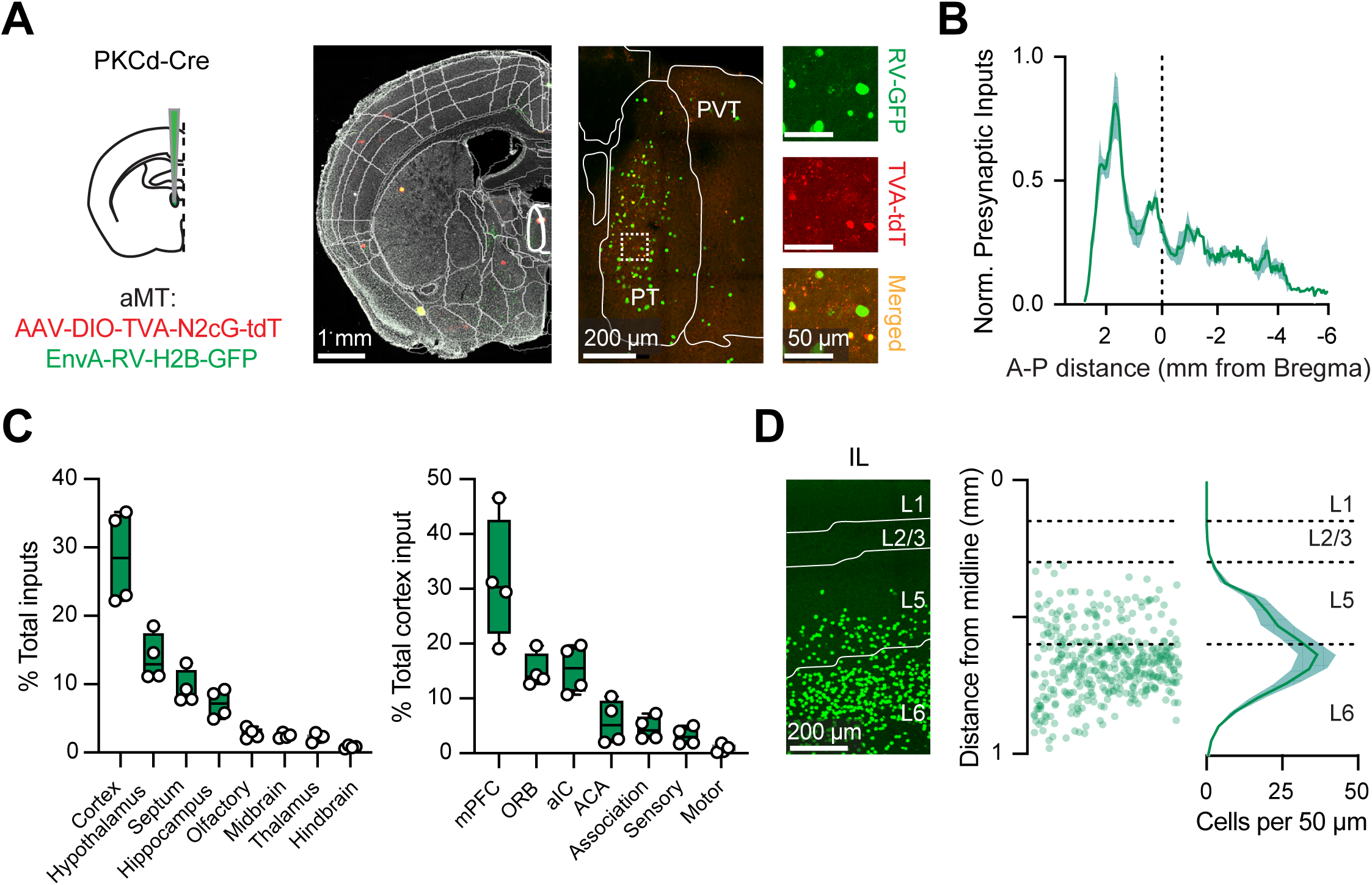
Identifying long-range inputs to PKCd+ cells in paratenial thalamus. **(A)** *Left*, Schematic of injections of AAV helper virus and pseudotyped RV into aMT of PKCd-Cre mice. *Right*, Images of coronal slice and magnification of aMT, showing GFP+ (green), tdTomato+ (red), and starter cells (yellow) in PT but not PVT. Scale bars = 1 mm, 200 µm, and 50 µm. **(B)** Summary of locations of presynaptic input cells to PKCd+ cells in PT across the anterior-posterior (A-P) axis of the brain (N = 4 mice). **(C)** *Left*, Summary of brain regions containing presynaptic input cells. *Right*, Summary of cortical presynaptic input cells. **(D)** *Left*, Image of corticothalamic (CT) input cells to PKCd+ PT cells in IL. *Middle*, Overlay of the position of all CT cells as a function of distance to the midline. *Right*, Summary of distribution of CT cells across cortical layers in 50 µm bins. Scale bar = 200 µm. Box and whisker plots represent median and minimum to maximum. Cell counts across cortical layers or A-P coordinates are presented as means ± SEM. *(see also Figure S4)*

### Prefrontal inputs selectively innervate the paratenial thalamus

Having identified mPFC as a major input to PT, we next examined the connections from corticothalamic (CT) cells in more detail. We first injected AAV-hSyn-eYFP into the mPFC of wild-type animals, waited for expression and transport, prepared coronal slices, and aligned to the CCF (**Fig. 5A**). Strikingly, we found that mPFC inputs almost entirely innervated PT, with minimal labeling of neighboring PVT (normalized intensity: PVT = 0.3 ± 0.1, PT = 0.9 ± 0.1, *p* < 0.0001, N = 5 mice) (**Fig. 5B**). In principle, these connections can arise from either L5 or L6 CT cells in the mPFC, which are known to have different properties and send distinct connections to other parts of thalamus (Collins *et al*., 2018). We thus used transgenic mice that selectively label mPFC inputs from layer 5 (L5) or layer 6 (L6) corticothalamic cells. Using Syt6-Cre mice that label deep layers of mPFC (Vaasjo *et al*., 2022), we found that L6 CT inputs again primarily innervated PT, with minimal labeling of PVT (normalized intensity: PVT = 0.2 ± 0.0, PT = 0.9 ± 0.0, *p* < 0.0001, N = 5 mice) (**Fig. 5C**). Using Rbp4-Cre mice that label other layers of mPFC (Lui *et al*., 2021), we found that L5 CT inputs showed less overall labeling, but still a bias to PT (normalized intensity: PVT = 0.1 ± 0.1, PT = 0.3 ± 0.1, *p* = 0.07, N = 3 mice) (**Fig. 5D**). These anatomical findings indicate that mPFC primarily innervates PT, with minimal connections in the adjacent PVT.

**Figure 5:**
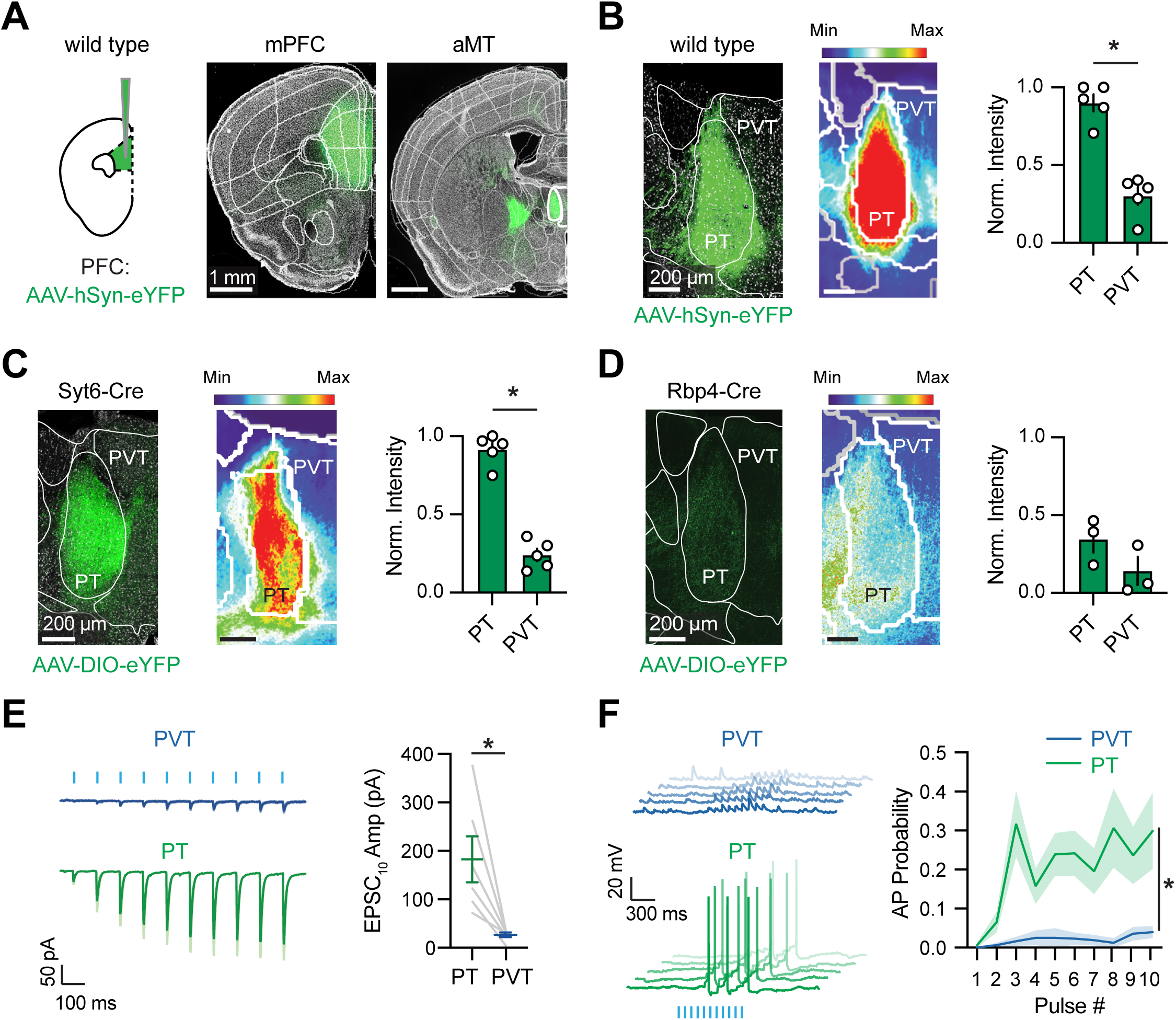
mPFC inputs preferentially innervate paratenial thalamus. (A) *Left*, Schematic of AAV-hSyn-YFP injection into the mPFC of wild-type mice. *Middle*, Representative image of injection site in mPFC. *Right*, Labeling of mPFC axons in aMT. Scale bars = 1 mm. (B) *Left*, Confocal image of mPFC axons in PT but not PVT. *Middle*, Average heat map from multiple slices (n = 3 sections per animal) and animals (N = 5 mice). *Right*, Summary of normalized fluorescence intensity in PT and adjacent PVT. Scale bars = 200 µm. (C) Similar to (B) for AAV-DIO-YFP injection into mPFC of Syt6-Cre mice. (D) Similar to (B) for AAV-DIO-YFP injection into mPFC of Rbp4-Cre mice. (E) *Left*, Average Syt6+ cell-evoked EPCSs recorded at E_GABA_ (-60 mV) in TC cells in PT (bottom, green) and PVT (top, blue). Scale bars = 50 pA x 100 ms. *Right,* Summary of EPSC_10_ amplitudes at TC cells in PT and PVT (n = 6 pairs, N = 4 mice). (F) *Left,* Syt6+ cell-evoked AP firing at TC cells in PT (bottom, green), and subthreshold EPSP in PVT (top, blue), with multiple traces overlaid. Scale bars = 20 mV x 100 ms. *Right*, Summary of AP probability as a function of pulse number (n = 8 pairs, N = 6 mice). Values are means ± SEM. * = p < 0.05. *(see also Figure S5)*

Previous studies demonstrate that mPFC communicates in tight reciprocal loops with other higher-order thalamic nuclei, including MD and VM thalamus (Collins *et al*., 2018; Halassa & Sherman, 2019). Therefore, we next asked if TC cells in PT are also contacted and driven to fire by mPFC inputs as part of a reciprocal loop between these areas. We combined similar injections described above, now injecting AAV-DIO-ChR2-YFP and AAVrg-H2B-tdTomato into mPFC of Syt6-Cre mice. After waiting for expression and transport, we made whole-cell recordings of pairs of TC cells residing in PT and PVT. To stimulate mPFC inputs, we used 10 Hz trains of blue light pulses (10 pulses at 10 Hz, 2 ms duration, 470 nm, and 4-15 mW power). In voltage-clamp recordings, we found that mPFC inputs evoked prominent EPSCs at TC cells in PT but only minimal responses at TC cells in the adjacent PVT (EPSC_10_: PVT = 27.1 ± 4.8 pA, PT = 182.6 ± 47.4 pA, *p* = 0.01, n = 6 pairs, N = 5 mice) (**Fig. 5E & S5A**). Responses built over stimulus trains, indicating presynaptic facilitation, consistent with L6 CT inputs to other thalamic nuclei (PPR_10_: PVT = 9.5 ± 1.5, PT = 8.8 ± 1.5) (**Fig. S5B & S5C**). In separate current-clamp recordings, we also found that mPFC inputs were sufficient to fire action potentials at TC cells in PT but not PVT (AP probability at pulse 10: PVT = 0.04 ± 0.01, PT = 0.3 ± 0.09, *p* = 0.02, n = 8 pairs, N = 6 mice) (**Fig. 5F**). These results show mPFC inputs preferentially contact TC cells in PT over PVT, consistent with our anatomy and suggesting a closed reciprocal loop between mPFC and PT.

### Prefrontal inputs preferentially contact subcortically projecting cells

Our results indicate that the mPFC makes connections onto TC cells in PT, forming a reciprocal loop between these structures. Given that the cortex- and subcortex-projecting populations within PT are non-overlapping, we next asked if mPFC also innervates the TS and TA cells that project to the NAc and BLA, respectively. In this case, we injected AAV-DIO-ChR2 and a retrograde tracer into the mPFC, along with another retrograde tracer into either NAc (N = 6 mice) or BLA (N = 7 mice) in Syt6-Cre mice. In voltage-clamp recordings, we found that mPFC inputs also made facilitating connections onto both TS and TA cells (**Fig. 6A**). However, in recordings from pairs of TC cells and neighboring TS or TA cells, we found responses were much larger at subcortically projecting cells (EPSC5: TC = 153.4 ± 41.9 pA, TS = 492 ± 114.8 pA, p = 0.01, n = 7 pairs; TC = 174.5 ± 95.11 pA, TA = 384.7 ± 111.7 pA, p = 0.002, n = 7 pairs) (**Fig. 6B & S6**). Biased inputs were similar for TS and TA cells (EPSC ratio: TS/TC = 3.5, 95% CI = 1.5-7.8, TA/TC = 3.3, 95% CI = 1.6-6.7, p = 0.84) (**Fig. 6C**), with similar short-term facilitation (**Fig. 6D**). In separate current-clamp recordings, we found that mPFC inputs also drive activity in TS and TA cells much more effectively than at TC cells (AP probability: TC = 0.1 ± 0.1, TS = 0.7 ± 0.1, p = 0.008, n = 7 TS/TC pairs, N = 7 mice; TC = 0.2 ± 0.1, TA = 0.8 ± 0.04, p = 0.001, n = 7 TA/TC pairs, N = 6 mice) (**Fig. 6E & 6F**). Synaptically evoked firing built during trains, allowing mPFC inputs to effectively engage these different cell populations. These findings indicate that while mPFC makes reciprocal connections with TC cells, it even more effectively engages subcortically projecting cells.

**Figure 6:**
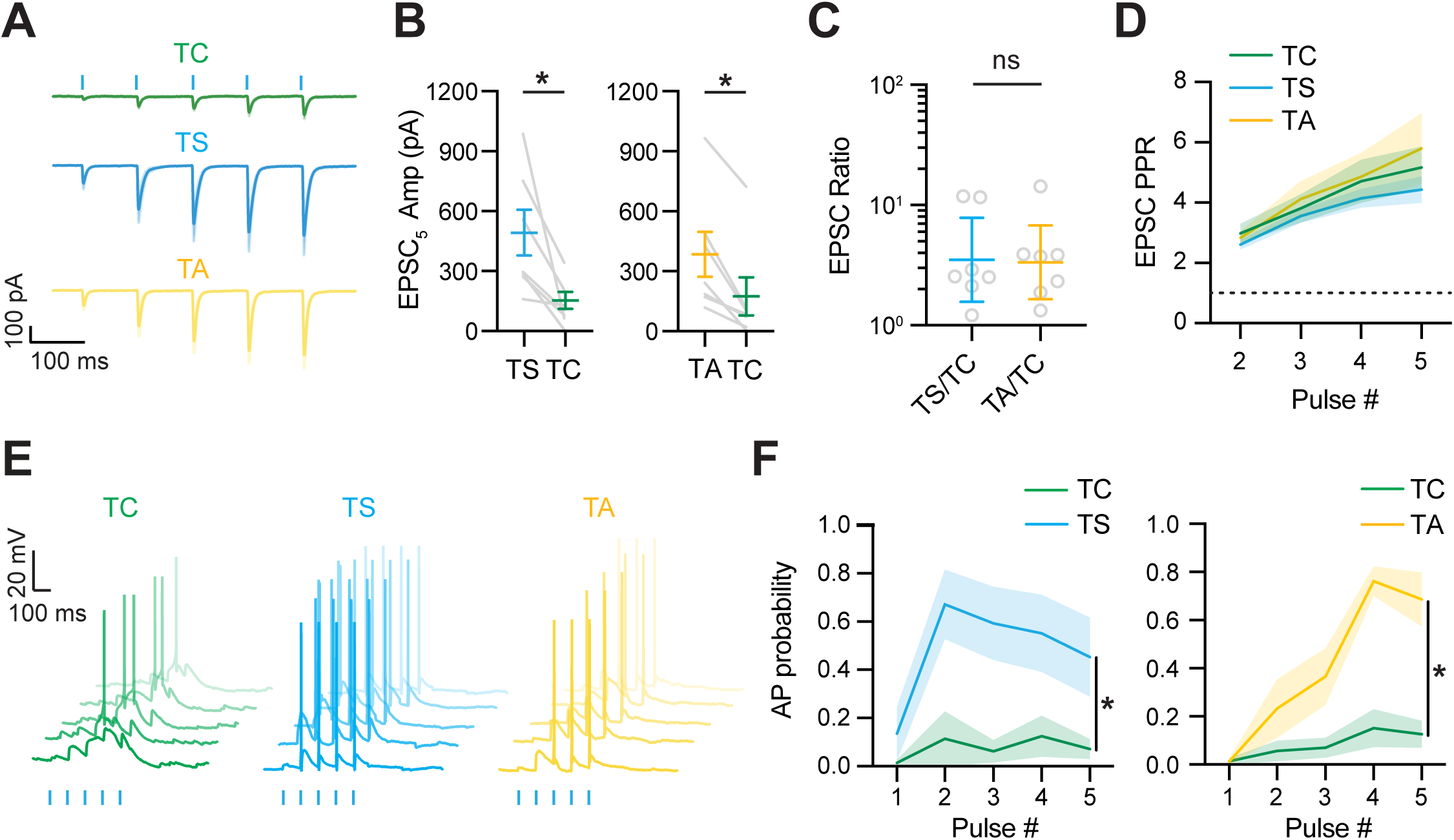
mPFC inputs are stronger onto subcortically projecting cells. (A) Average Syt6+ cell-evoked EPCSs recorded at E_GABA_ (-60 mV) in TC (green), TS (blue), and TA (yellow) cells in PT. Scale bars = 100 pA x 100 ms. (B) Summary of EPSC_5_ amplitudes in pairs of TC and TS cells (left) (n = 7 pairs, N = 6 mice), and pairs of TC and TA cells (right) (n = 7 pairs, N = 6 mice) in PT. (C) Summary of EPSC ratio for TC vs. TS and TC vs. TS in PT. (D) Summary of PPR as a function of stimulation pulse number. Dotted line indicates PPR = 1. (E) Syt6+ cell-evoked AP firing in TC, TS, and TA cells in PT with multiple traces overlaid. Scale bars = 20 mV x 100 ms. (F) Summary of Syt6+ cell-evoked AP probability as a function of stimulation pulse number for TC vs TS (left) (n = 7 pairs, N = 7 mice), and TC vs TA (right) (n = 7 pairs, N = 6 mice). Values are means ± SEM or geometric means ± 95% CI. * = p < 0.05. *(see also Figure S6)*

### Polysynaptic circuits link mPFC to NAc and BLA via paratenial thalamus

Together, our findings suggest that CT cells in the mPFC can indirectly access both the NAc and BLA via PT. However, testing this idea is complicated by the lack of a specific genetic marker for TS and TA cells in this nucleus. As an alternative strategy, we took advantage of our finding that mPFC inputs selectively target PT over PVT. We injected AAV1-Cre in mPFC, which at high titer travels anterogradely and expresses Cre in postsynaptic cells (Zingg *et al*., 2017). In the same animals, we also injected AAV-DIO-ChR2-eYFP into aMT, which we found selectively labeled PT over PVT (**Fig. 7A**). In this case, we observed green axons in both cortical and subcortical brain regions, including cortex, striatum, and amygdala (**Fig. 7B & 7C**). In the striatum, we found that labeling was concentrated in the medial core and shell of the NAc (**Fig 7B**), similar to previous work on PVT (Gimenez-Amaya *et al*., 1995; Vertes & Hoover, 2008; Zhu *et al*., 2016; Shima *et al*., 2023). In the amygdala, we found labeling was mainly in the medial BLA (**Fig. 7B**), but sparse in the central nuclei. Lastly, to examine functional connection, we prepared slices and recorded from pyramidal cells in IL, MSNs in NAc, and pyramidal cells in BLA. In voltage-clamp, we found that PT inputs evoked prominent EPSCs in each of these brain regions (EPSC_1_: IL = 263.5 ± 47.5 pA, n = 8 cells, NAc = 213.2 ± 33.5 pA, n = 6 cells, BLA = 235.7 ± 75.61 pA, n = 7 cells, N = 8 mice) (**Fig. 7D**), all of which showed short-term depression (PPR_5_: IL = 0.5 ± 0.1, NAc = 0.6 ± 0.07, BLA = 0.7 ± 0.1) (**Fig. S7A)**. PT inputs also evoked feed-forward inhibition, suggesting engagement of local inhibitory circuits (**Fig. S6B-D).** Lastly, in separate current-clamp recordings, we found that PT inputs could evoke transient action potential firing in each of these target cells, with highest firing rates earlier in the stimulation trains (AP probability at pulse 1: IL = 0.7 ± 0.1, NAc = 0.5 ± 0.1, BLA = 0.5 ± 0.1) (**Fig. 7E**). Together, these findings indicate that TC, TS and TA cells in PT make functional, potent connections onto both cortical and subcortical targets.

**Figure 7:**
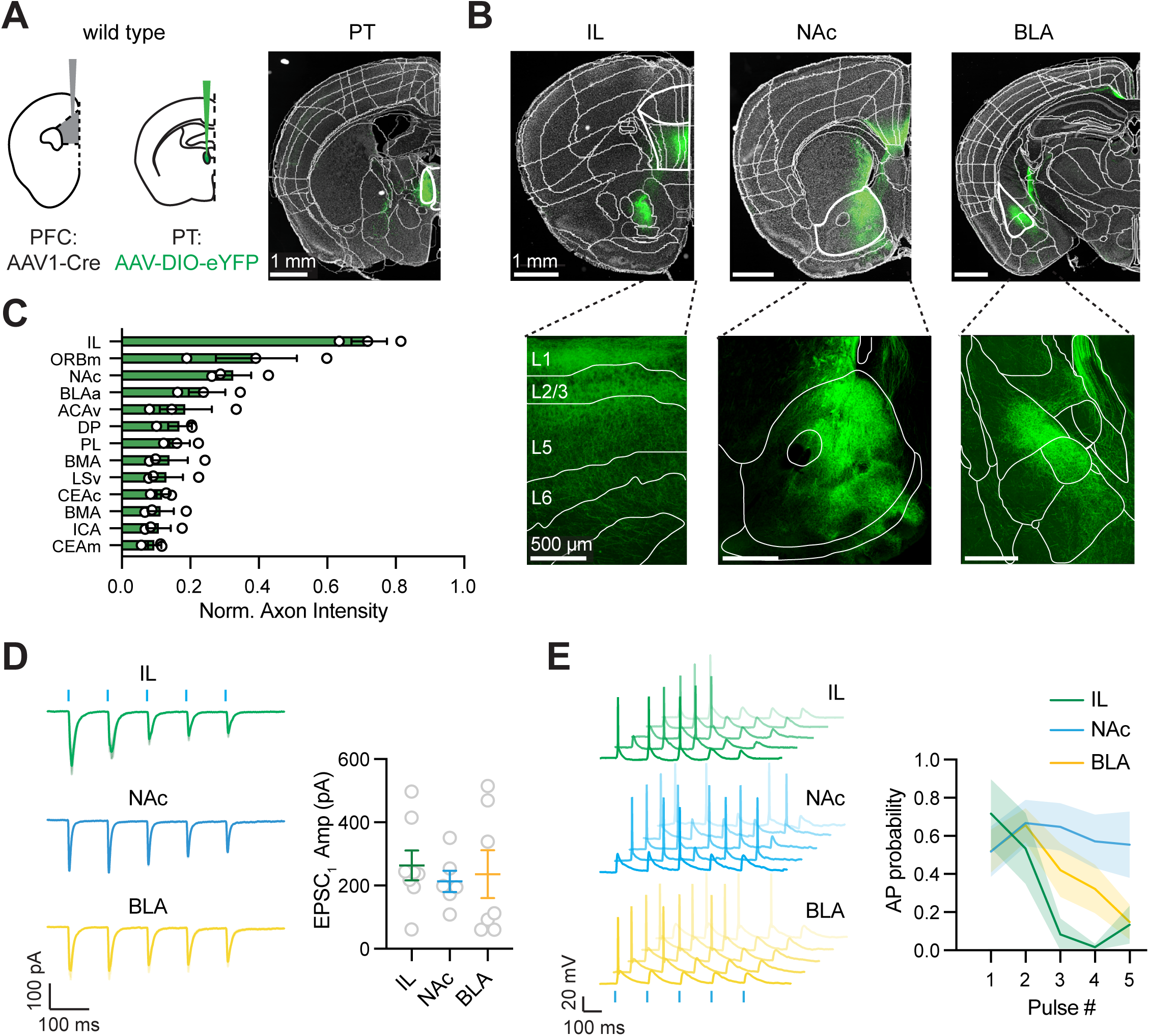
Paratenial thalamus routes cortical signals to limbic areas. **(A)** *Left*, Schematic of injections of AAV1-hSyn-Cre into the mPFC and AAV-DIO-eYFP into PT of wild-type mice. *Right*, Image of CCF-aligned coronal section, showing viral expression in PT. **(B)** *Top,* Coronal slices showing PT axons in IL (left), NAc (middle), and BLA (right). *Bottom,* Magnification of IL, NAc, and BLA. Scale bars = 1 mm (top) and 500 µm (bottom). **(C)** Summary of PT axonal fluorescence intensity across areas (N = 3 mice). **(D)** *Left,* Average trans-synaptically labeled PT-evoked EPSCs at IL L2/3 pyramidal cells (top; green), NAc MSNs (middle; blue), and BLA pyramidal cells (bottom; yellow). Scale bars = 100 pA x 100 ms. *Right,* Summary of EPSC_1_ amplitudes in IL, NAc, and BLA (n = 7-8 cells per area, N = 5 mice). **(E)** *Left,* Trans-synaptically labeled PT-evoked AP firing in IL L2/3 pyramidal neurons (top), NAc MSN (middle), and BLA pyramidal cells (bottom), with multiple traces overlaid. Scale bars = 20 mV x 100 ms. *Right,* Summary of evoked AP probability as a function of stimulation pulse number (n = 8-9 cells per group, N = 5-6 mice). Values are means ± SEM. *(see also Figure S7)*

## DISCUSSION

Our findings indicate that paratenial thalamus (PT) and paraventricular thalamus (PVT) are genetically, anatomically, and functionally distinct nuclei in anterior midline thalamus (aMT). PKCd is a unique genetic marker for thalamocortical (TC) neurons in PT, while thalamostriatal (TS) and thalamoamygdala (TA) neurons represent a smaller and distinct population of subcortically projecting neurons in PT and PVT. PKCd+ TC cells primarily project to superficial layers of infralimbic (IL) cortex, where their axons overlap with the endogenous PKCd in thalamo-recipient layers and drive transient activity in L2/3 pyramidal neurons in IL. Reciprocal corticothalamic projections from deep layers of the mPFC are also heavily localized in PT over PVT, where they target and strongly drive activity in subcortex-projecting neurons. Together, our data demonstrate PT as a key node in the cortico-thalamo-limbic network, enabling mPFC cortical signals to be routed to limbic areas, including the nucleus accumbens (NAc) and basolateral amygdala (BLA).

The borders between PT and PVT have been difficult to delineate, and historically drawn using peptidergic staining or cytoarchitecture (Levey *et al*., 1987; Groenewegen & Berendse, 1994; Kirouac, 2015), such that nuclei are often grouped together. Recent reports highlight the significant genetic diversity in PVT along its anterior-posterior axis (Gao *et al*., 2020; Gao *et al*., 2023; Shima *et al*., 2023). However, to our knowledge, there have been no equivalent studies on the molecular signatures of PT. Previously reported PVT-specific markers, including *ntrk1* (Shima et al., 2023) and *nts* (Li et al., 2022), are not expressed in PT. *Calb2* is often associated with PVT (Matyas *et al*., 2018; Viena *et al*., 2021; Biro *et al*., 2025), but we find it is ubiquitously expressed in dorsal aMT, including both PT and PVT. Moreover, *calb2* is not specific for any of the three main projection populations we studied in either PT or PVT. On the other hand, *prkcd* is uniquely enriched in PT, approximately 10-fold higher than PVT. Thus, our data indicates that PT and PVT are genetically distinct nuclei, potentially with distinct connectivity and functional roles.

Similarly, our retrograde anatomy data reveal differential distribution of TC, TS, and TA neurons in PT and PVT. Mirroring the distribution of PKCd+ neurons, TC neurons are more abundant in PT, in accordance with previous tracing studies in macaques (Hsu & Price, 2007), cats (Room *et al*., 1985), and rats (Vertes & Hoover, 2008). Subcortex-projecting neurons, including TS and TA neurons, are more abundant in PVT compared to PT. Interestingly, within PT, TC neurons are co-labeled with PKCd, with minimal overlap with the TS and TA populations. While these thalamic cells are historically categorized as a homogenous population that projects multiple areas (Groenewegen & Berendse, 1994; Matyas *et al*., 2018), our data indicates that there are at least two distinct streams of PT outputs, one targeting the mPFC and the other projecting to subcortical limbic structures, including NAc and BLA. All of these projection-specific PT populations appear to share similar intrinsic physiological properties, including tonic firing at depolarized potentials and burst firing following hyperpolarization, and dendritic morphology (Llinás & Jahnsen, 1982).

Thalamocortical projections from high-order thalamic nuclei arrive in multiple superficial layers of the mPFC, including layers 1 and 2/3 (Castro-Alamancos & Connors, 1997; Jones, 1998). Using PKCd-Cre mice allowed us to restrict viral expression to PT and avoid PVT, which has previously been challenging due to the small size and close proximity of these nuclei. In agreement with our retrograde tracing data, PKCd+ TC cell axons are most densely labeled in IL, with minimal axons in NAc and BLA. Moreover, PKCd+ TC cell axons in IL resemble those of mediodorsal (MD) thalamus in prelimbic (PL), terminating in superficial layers L1 and L2/3 (Collins *et al*., 2018; Anastasiades *et al*., 2021). Similarly, PKCd+ TC synapses display pronounced short-term depression and transiently drive activity in L2/3 pyramidal neurons. PKCd+ TC inputs also evoke feed-forward inhibition in IL, indicating they engage local inhibitory networks. Recent work indicates MD inputs activate PV+ interneurons in anterior cingulate cortex (ACC) (Delevich et al., 2015), but CCK+ interneurons in PL (Kamalova et al., 2024). In future studies, it will be interesting to further characterize the influence of PKCd+ TC synapses on specific microcircuits in IL.

Our anatomy indicates that PKCd expression also closely overlaps with TC axons in mPFC and strongly labeled other cortical areas. For example, PKCd is robustly expressed in thalamo-cortical recipient layers in visual (L1 and L4), somatosensory (L4 barrels), and motor cortices (L3), suggesting it may serve as a common marker for thalamocortical connections. Intriguingly, PKC is also a key effector in Gq-mediated signaling pathway (de Jong & Verhage, 2009), suggesting it may play a role in regulation of TC synapses. In the future, it will thus be interesting to assess how PKCd influences both modulation and plasticity at TC connections in mPFC and elsewhere.

The aMT is often considered an integrative hub that receives convergent inputs from limbic areas involved in arousal and reward. Our rabies-based tracing experiments of presynaptic inputs to PKC+ cells confirmed that PT samples inputs from a wide array of cortical and subcortical areas for subsequent routing to cortex and subcortex. The largest subcortical inputs come from a collection of hypothalamic nuclei, many of which do not send direct connections to IL (Schaeuble & Myers, 2022). PKCd+ cells may therefore act as a relay center of interoceptive and homeostatic information to IL for top-down control of behavior. Interestingly, ventral hippocampus (vHIPP) sends abundant inputs to PKCd+ cells and is also directly connected with IL (Liu & Carter, 2018). Our results therefore suggest that PT thalamus may serve as an additional waystation for the routing of hippocampus signaling to the prefrontal cortex, similar to the nucleus reuniens (Dolleman-van der Weel *et al*., 2019). In future studies, it will be interesting to compare the connectivity and function of these direct and indirect hippocampal-prefrontal connections.

While subcortical inputs are present, the major input to the PT is from frontal cortex, with densest presynaptic cell labeling in IL. Strikingly, anterograde labeling of corticothalamic (CT) inputs from IL predominantly labels PT, with minimal labeling of nearby PVT. Similarly, CT inputs preferentially engaged TC cells in PT, with minimal functional connections onto TC cells in PVT. These inputs primarily derive from L6 CT cells, which make facilitating synapses that can robustly drive TC cells, with relatively little input from L5 CT cells. These findings are reminiscent of a similar closed reciprocal loop that links the mPFC with both MD and VM thalamus (Collins *et al*., 2018), in this case with both TC cells and CT inputs restricted to PT. However, our findings stand in contrast to the prevailing view that the mPFC strongly and reciprocally engages PVT (Otis *et al*., 2017; Gao *et al*., 2020; Aquino-Miranda *et al*., 2024; Ma *et al*., 2024), instead indicating these interactions are confined to PT. One possible explanation is the historical difficulty in distinguishing PT from PVT, as our findings indicate that the commonly used marker Calb2 labels both of these closely spaced nuclei. Consequently, some anatomical tracing and electrophysiological recordings could have been inadvertently performed in PT instead of PVT.

Most studies on cortico-thalamo-cortical interactions have focused on reciprocal connections between thalamus and cortex, including somatosensory (Reichova & Sherman, 2004; Theyel *et al*., 2010; Crandall *et al*., 2015), visual (Andolina *et al*., 2007; Olsen *et al*., 2012; Born *et al*., 2021), auditory (Bartlett & Smith, 2002; Ibrahim *et al*., 2021) and higher-order systems (Schmitt *et al*., 2017; Collins *et al*., 2018). In addition to TC cells, PT also contains intermingled TS and TA cells that project to subcortical brain regions. These three populations of cells are also found in PVT, but less so in other higher-order nuclei like MD and VM. Facilitating mPFC inputs evoke robust firing of TC, TS, and TA cells in PT, indicating they can drive sustained activity. However, responses are largest at TS and TA cells, indicating mPFC preferentially drives subcortically projecting cells. Therefore, the mPFC inputs can simultaneously regulate activity of multiple cell types in PT, including those that constitute closed reciprocal loops as well as broadcast networks. Together, our data suggests that PT is optimized for routing prefrontal cortical signals to the limbic system, in contrast to MD thalamus, which forms closed cortico-thalamo-cortical loops that amplify and sustain cortical activity (Schmitt *et al*., 2017). More broadly, our findings show how mPFC connections with different thalamic nuclei may perform distinct computational and functional roles.

Our transsynaptic labeling experiments using AAV1 capitalized on our finding that mPFC axons are heavily localized to PT over PVT. This viral strategy proved useful not only for isolating PT, but also for labeling PT cells that project to NAc and BLA. We observed that mPFC-recipient PT cells project densely to the dorsomedial shell and core of the NAc, as well as the medial wall of the BLA. PT excitatory inputs to IL, NAc and BLA display short-term depression, as seen in other thalamocortical (Cruikshank *et al*., 2010; Cruikshank *et al*., 2012; Kamalova *et al*., 2024), thalamostriatal (Ding *et al*., 2008), and thalamoamygdala synapses (Szinyei *et al*., 2000), and are sufficient to drive firing in these areas. In each case, PT inputs can also drive feed-forward inhibition, confirming that they recruit local interneurons. In future studies, it will be interesting to determine which principal cells and interneurons are engaged by PT inputs in NAc and BLA.

More broadly, these experiments suggest multiple routes through which cortical signals can access limbic structures via PT. The mPFC has direct connections with the NAc and BLA, which arise from pyramidal cells in layers 2/3 and 5 (Anastasiades *et al*., 2019). In contrast, mPFC makes connections to PT via separate CT cells located in layers 5 and 6. These projection-specific populations belong to two classes of cortical neurons with distinct genetic, physiological, and morphological properties (Gabbott *et al*., 2005; Oberlaender *et al*., 2012; Anastasiades & Carter, 2021; Yao *et al*., 2021). By driving activity in subcortex-projecting PT cells, mPFC signals can thus be broadcast to multiple subcortical structures. In the dorsal striatum, cortical and thalamic synapses can target different patch and matrix compartments (Fujiyama *et al*., 2006), and have different synaptic properties (Finch, 1996; Ding *et al*., 2008; Smeal *et al*., 2008). Similarly, cortical and thalamic inputs are thought to engage different cell types and local circuits in the amygdala (Chen *et al*., 2022; Li *et al*., 2022). It will be interesting to compare the neural circuits engaged by these direct cortico-limbic and indirect cortico-thalamo-limbic pathways in NAc and BLA.

In summary, our findings help to revise the canonical view of the aMT by identifying PT as a molecularly distinct nucleus that channels prefrontal output to limbic structures. Dorsal aMT circuits are involved in a multitude of behaviors and physiological states, including arousal (Schmitt *et al*., 2017; Gao *et al*., 2020), fear learning (Do-Monte *et al*., 2015), reward seeking (Otis *et al*., 2017), and social behaviors (Yamamuro *et al*., 2020), and are implicated in a host of neuropsychiatric disorders, including stress (Kooiker *et al*., 2023), pain (Tang *et al*., 2024), and substance use disorders (Zhu *et al*., 2016; Clark *et al*., 2017; Paniccia *et al*., 2024). Our anatomical and physiological characterizations of PT circuits lay a foundation for understanding PT regulation of the limbic system, including elucidating functional impact on emotional processing and motivated behaviors, with implications for related neuropsychiatric disorders.

## MATERIALS & METHODS

All experimental procedures were approved by the New York University Animal Welfare Committee. P28-P70 wild-type (JAX #000664), Syt6-Cre (RRID: MMRRC_037416-UCD), and Rbp4-Cre (RRID: MMRRC_037128-UCD) mice of both sexes were used on a C57BL/6J background. PKCd-Cre BAC transgenic mice were generated by GENSAT and generously provided by Eric Klann (Haubensak *et al*., 2010). All animals were held in reverse light cycle and recorded in the dark cycle. All experiments were replicated in at least 3 animals. No formal method for randomization was used, and experimenters were not blind to experimental groups. No pre-test analyses were used to estimate sample sizes. No data were excluded from final analyses.

### Stereotaxic injections

Mice were anesthetized with isoflurane (1-2%) and head fixed in a stereotax (Kopf Instruments). A small craniotomy was made over the injection site, at the following coordinates relative to bregma (mediolateral, dorsoventral, anteroposterior): mPFC = ±0.35, -2.5, +2.1 mm; PT = ±1.5, –3.9, +0.4 mm at a 18° angle; NAc = ±1.8, -4.3, +1.4 mm at a 14° angle; BLA = ±3.1, -5.1, -1.9 mm. Borosilicate pipettes with 5-10 µm tip diameters were backfilled with mineral oil. Injection volumes of 100 nl (into PT) or 150-300 nl (into mPFC, NAc, and BLA) of virus were pressure-injected using a Nanoject III (Drummond) with 30-45 second inter-injection intervals. For retrograde labeling, mice were injected with either G-deleted rabies virus RV-CVS-N2c(ΔG)-H2B-GFP (UNC Vector Core, titer = 3.13E+08 GC/ml), AAVrg-CAG-H2B-tdTomato (UNC Vector Core, NT-231075.1, titer = 2.18E+13 GC/ml), Alexa-647-tagged cholera toxin subunit B (CTB647, Thermo Fisher, C34778) or a 1:1:1 mixture of rabies virus, AAVrg, and CTB. For tracing of monosynaptic inputs to PKCd+ cells in PT, PKCd-Cre mice were first injected with a helper virus AAV1-hSyn-DIO-TVA66T-dTom-CVS-N2cG (UNC Vector Core, titer = 2.81E+13 GC/ml) in PT. Three weeks later, pseudotyped rabies EnVA-CVS-N2c(ΔG)-H2B-GFO (UNC Vector Core, titer = 1.0E+9 GC/ml) were injected into the same location. Anterograde Cre-dependent anterograde labelling was achieved using AAV1-EF1a-DIO-eYFP-WPRE.hGH (Addgene #27056-AAV1, titer = 2.5E+13 GC/ml) for anatomical tracing, and AAV1.EF1a.DIO.hChR2(H134R)-eYFP.WPRE.hGH (Addgene #20298-AAV1, titer = 2.3E+13 GC/ml) for optogenetic stimulation. For transsynaptic labeling of PT, AAV1-hSyn-Cre-WPRE.hGH (Addgene #105553-AAV1, titer = 1.9E+13 GC/ml) was injected into mPFC, and AAV1.EF1a.DIO.hChR2(H134R)-eYFP.WPRE.hGH was injected into PT. Following injections, the pipette was left in place for an additional 5-10 min before being slowly withdrawn from the brain. Animals received post-op analgesics (ketoprofen, 5 mg/kg, 3 days) and returned to their home cages, where they remained for 2-4 weeks before experiments.

### Slice preparation

Mice were deeply anesthetized with isoflurane and perfused intracardially with an ice-cold cutting solution containing (in mM): 65 sucrose, 76 NaCl, 25 NaHCO_3_, 1.4 NaH_2_PO_4_, 25 glucose, 2.5 KCl, 7 MgCl_2_, 0.4 Na-ascorbate, and 2 Na-pyruvate (bubbled with 95% O_2_/5% CO_2_). 300 µm coronal sections were cut in this solution and transferred to ACSF containing (in mM): 120 NaCl, 25 NaHCO_3_, 1.4 NaH_2_PO_4_, 21 glucose, 2.5 KCl, 2 CaCl_2_, 1 MgCl_2_, 0.4 Na-ascorbate, and 2 Na-pyruvate (bubbled with 95% O_2_/5% CO_2_). Slices were recovered for 30 min at 35°C and stored for at least 30 min at room temperature. All recordings were conducted at 30-32°C.

### Slice electrophysiology

Whole-cell recordings were made from neurons using infrared-differential interference contrast as described previously (Collins *et al*., 2018; Anastasiades *et al*., 2021; Kamalova *et al*., 2024). In the PT and PVT, retrogradely labeled neurons were identified by virally or CTB-expressed fluorophores. Sequential pairs of TC cells in PT vs. PVT, TC vs. TS cells in PT, or TC vs. TA cells in PT, were recorded in alternated orders to eliminate biases. For voltage-clamp experiments, borosilicate pipettes (3-5 MΩ) were filled with the following (in mM): 135 Cs-gluconate, 10 HEPES, 10 Na-phosphocreatine, 4 Mg_2_-ATP, 0.4 NaGTP, 10 TEA, 2 QX-314, and 10 EGTA, pH 7.3 with CsOH (290-295 mOsm). For current-clamp recordings, borosilicate pipettes (3-5 MΩ) were filled with the following (in mM): 135 K-gluconate, 7 KCl, 10 HEPES, 10 Na-phosphocreatine, 4 Mg_2_-ATP, 0.4 NaGTP, and 0.5 EGTA, pH 7.3 with KOH (290-295 mOsm). In some cases, 30 µM Alexa Fluor-594 (Thermo Fisher, A10438) was added to visualize morphology with two-photon microscopy. In all optogenetic stimulation experiments, 10 µM (R)-CPP was used to block NMDA receptors (Tocris Bioscience #0247), in order to be able to detect inhibition as needed. All reagents were purchased from Sigma or Tocris Bioscience.

Slice electrophysiology data were collected with a Multiclamp 700B amplifier (Axon Instruments) and National Instruments boards using custom software in MATLAB (MathWorks). Signals were sampled at 10 kHz and filtered at either 5 kHz for current-clamp recordings or 2 kHz for voltage-clamp recordings. Series resistance was 10-25 MΩ and not compensated.

### Slice optogenetics

Glutamate release was triggered by activating channelrhodopsin-2 (ChR2) present in presynaptic terminals of mPFC inputs to the thalamus (Collins et al., 2018). Channelrhodopsin-2 was activated using 2 ms pulses of 473 nm light from a blue light-emitting diode (LED; 473 nm; Thorlabs) delivered via a 10X 0.3 NA objective centered over the recorded neuron (Olympus), with a power range of 2-15 mW, routinely measured and calibrated at the back focal plane of the objective.

### Two-photon microscopy

Two-photon imaging was performed using a custom microscope and software, as previously described (Chalifoux and Carter, 2010). A titanium:sapphire laser (Coherent) tuned to 810 nm was used to excite Alexa Fluor 594 to image dendrite morphology. Z-stack images were subsequently acquired with a 60x 1.0 NA objective (Olympus) and processed with NIH ImageJ.

### Histology and immunohistochemistry

Mice were deeply anesthetized with isoflurane and perfused intracardially with 0.01 M PBS followed by 4% PFA. Brains were stored in 4% PFA overnight at 4°C before being washed three times in 0.01 M PBS. Slices were cut on a VT-1000S vibratome (Leica) at 60 µm thickness. For immunohistochemistry, slices were washed once in PBS (0.01 M), once in PBS-T (0.2% Triton-X100). Non-specific binding was blocked with PBS-T with 1% w/v bovine serum albumin (BSA) and 2% normal goat serum (Abcam #ab7481) for 1 hr at room temperature. Slices were incubated in a mixture of primary antibodies (mouse anti-calretinin, MA5-47457, Thermo Fisher, 1:2000, rabbit anti-protein kinase c delta, ab182126, Abcam, 1:2000) at 4°C overnight. After washing 4x in PBS, slices were incubated with secondary antibodies (goat anti-mouse IgG H&L Alexa 488-conjugated, A28175, Abcam, 1:500; goat anti-rabbit Alexa 647-conjugated, A21244, Abcam, 1:500) for 1 hr at room temperature. Slices were washed a further 3x in PBS before being mounted on gel-coated glass slides and cover-slipped with VectaShield with DAPI (Vector Labs). Fluorescent images were taken with an Olympus VS120 microscope, using 10X 0.25 NA objective (Olympus) or a Leica TCS SP8 confocal microscope, using 20X 0.75 NA objective (Olympus).

### Fluorescence in situ hybridization (RNAscope)

Mice were anesthetized with isofluorane and then perfused intracardially with chilled 0.01 M PBS. Brains were immediately submerged in cold isopentane on dry ice after dissection and stored in an airtight container at −80°C until sectioning. Sectioning was performed on a cryostat at −13°C, and 10 µm slices were mounted on Superfrost Plus microscope slides (Fisher Scientific) and stored at −80°C until staining. We followed a standardized RNAscope protocol for flash-frozen tissue from ACDBio, using Mm-nts (420441), Mm-nos1 (437651), Mm-calb2-C2 (313641-C2), Mm-ntrk1-C2 (435791-C2), and Mm-prkcd_C3 (441791-C3) probes. Slides were mounted using ProLong Gold antifade reagent with DAPI (Invitrogen) before being covered under coverslips.

### Data analysis

Off-line analysis of slice physiology recordings was performed using Igor Pro (WaveMetrics). For current-clamp recordings, resting membrane potential (Vrest) was measured at break-in. Input resistance (Rin) was measured using the steady-state response to a 500 ms current steps of –25 or –50 pA. Voltage sags indicating h-current were calculated by taking the minimum voltage in the first 200 ms, subtracting the average voltage over the final 50 ms, and dividing by the steady-state value. Optically evoked firing probabilities were calculated as the numbers of the action potentials within a 100 ms window divided by the numbers of trials. For voltage-clamp recordings, PSC amplitudes were measured as the average value across 1 ms around the peak response.

Coronal brain sections were registered to the Allen Institute’s Common Coordinate Framework (CCF) (Wang *et al*., 2020) using the DAPI channel. Retrogradely labeled, immunohistochemically labeled, or FISH-labeled cells were automatically counted using NeuroInfo software (MBF Bioscience). For analysis of co-localization of PKCd and Calb2 in retrogradely-labeled neurons, cells were manually counted in ImageJ. Thalamic axon distributions across the brains were quantified using NeuroInfo software. The average fluorescence intensity for each region was subtracted from within-slice background fluorescence and normalized to the peak fluorescence intensity across all measured areas.

For analysis of two-photon image stacks, all visible dendrites of each neuron were reconstructed in Neurolucida (MBF Bioscience). For Sholl analysis, 10 µm concentric circles were centered on the soma, and dendritic intersections and total dendritic lengths were calculated.

### Statistical Analysis

N values are reported within the Results and Supplemental Figure legends as the number of recorded cells (n) and animals (N) (for physiology) or number of animals (N) (for anatomy). Summary data are reported in the text and figures as arithmetic mean ± SEM. Ratio data are displayed in figures on logarithmic axes and reported as geometric mean ± 95% confidence interval (CI). In some graphs with three or more traces, SEM waves are omitted for clarity. Statistical analyses were performed using Prism 9.0 (GraphPad Software). Comparisons between unpaired data were performed using two-tailed Student’s t-tests. Comparisons between data recorded in pairs were performed using two-tailed paired t-tests. Comparisons of more than two groups were performed using two-tailed one-way ANOVA with Tukey’s multiple comparisons tests. Comparisons between groups across a range of variables (FI curves, PSC amplitudes and action potential probability per stimulation pulse) were made using repeated-measures two-way ANOVA with Sidak’s multiple comparison test. For all comparisons, no assumptions were made regarding data distributions, and significance was defined as p < 0.05.

## Acknowledgements

We thank members of the Carter lab, Paul Anastasiades, David Collins, and members of the U19 thalamus team for helpful discussions and comments on the manuscript. This work was supported by NIH R01 MH085974 (AGC) and NIH U19 NS123714 (AGC). The authors declare no competing interests.

---

**Figure S1:**
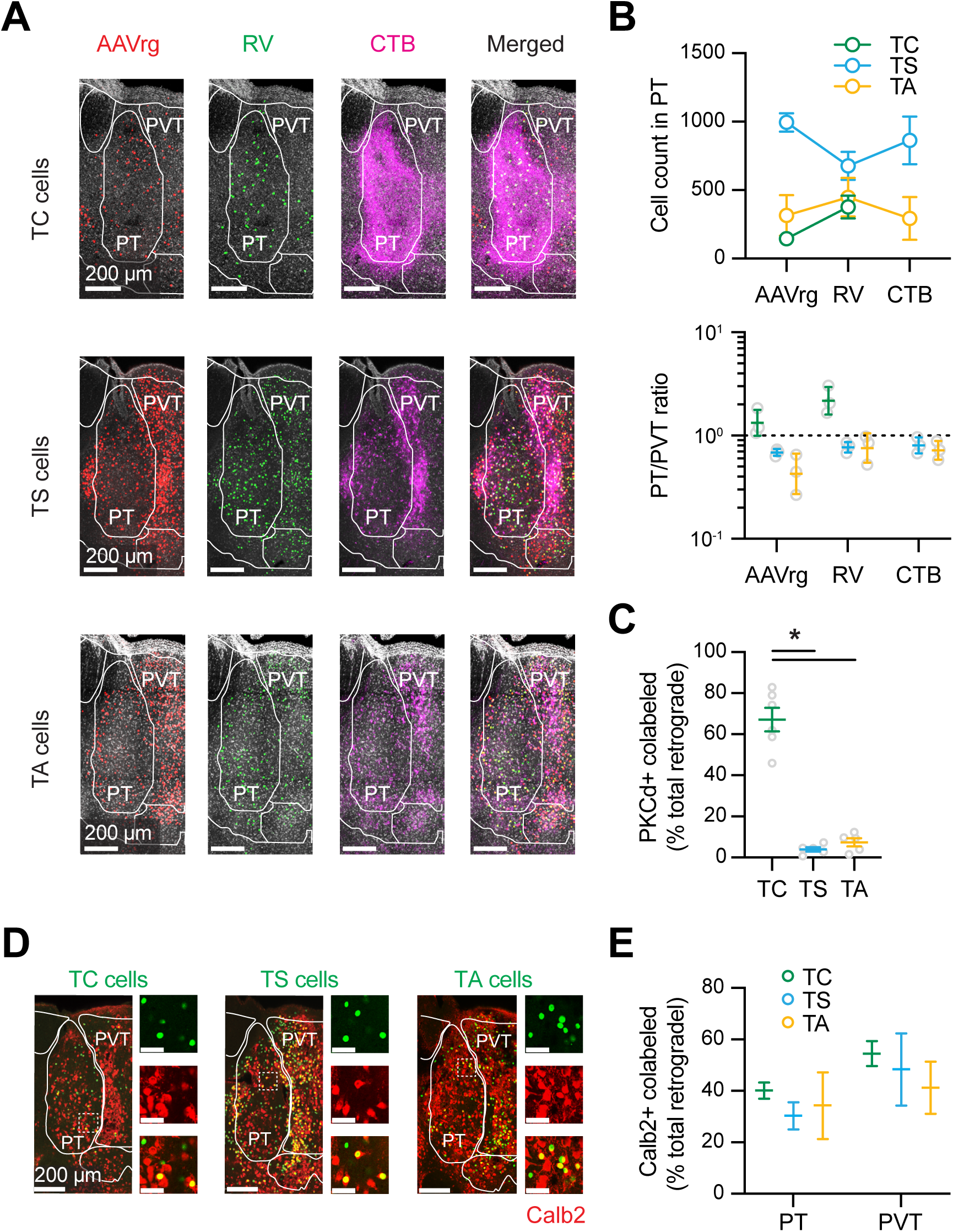
Further characterization of projection neurons in aMT. **(A)** Confocal images of TC cells, TS cells, and TA cells, labeled with co-injections of AAVrg-CAG-tdTomato (red), RV-GFP (green), and CTB647 (magenta) into mPFC (top), NAc (middle), and BLA (bottom), respectively. Note that strong labeling of CTB647 in corticothalamic axons from mPFC (see Figure 5) precludes quantification of TC cells in PT. Scale bars = 200 µm. **(B)** *Top*, Summary of retrogradely labeled cells across the 3 different methods. *Bottom,* Summary of PT/PVT ratio of retrogradely labeled cells (N = 3 mice per group). **(C)** Summary of PKCd+ co-labeled cells as a percentage of total retrogradely labeled cells (N = 5 mice per group). **(D)** Confocal images of TC, TS, and TA cells co-labeled with Calb2 in aMT, with magnified images showing GFP expression (top), Calb2 IHC (middle), and merge (bottom). Scale bar = 200 µm or 50 µm. **(E)** Summary of Calb2+ TC, TS, and TA cells in PT and PVT (N = 3 mice per group). Values are mean ± SEM or geometric means ± 95% CI (panel B). * = *p* < 0.05 *(Related to Figure 2)*

**Figure S2:**
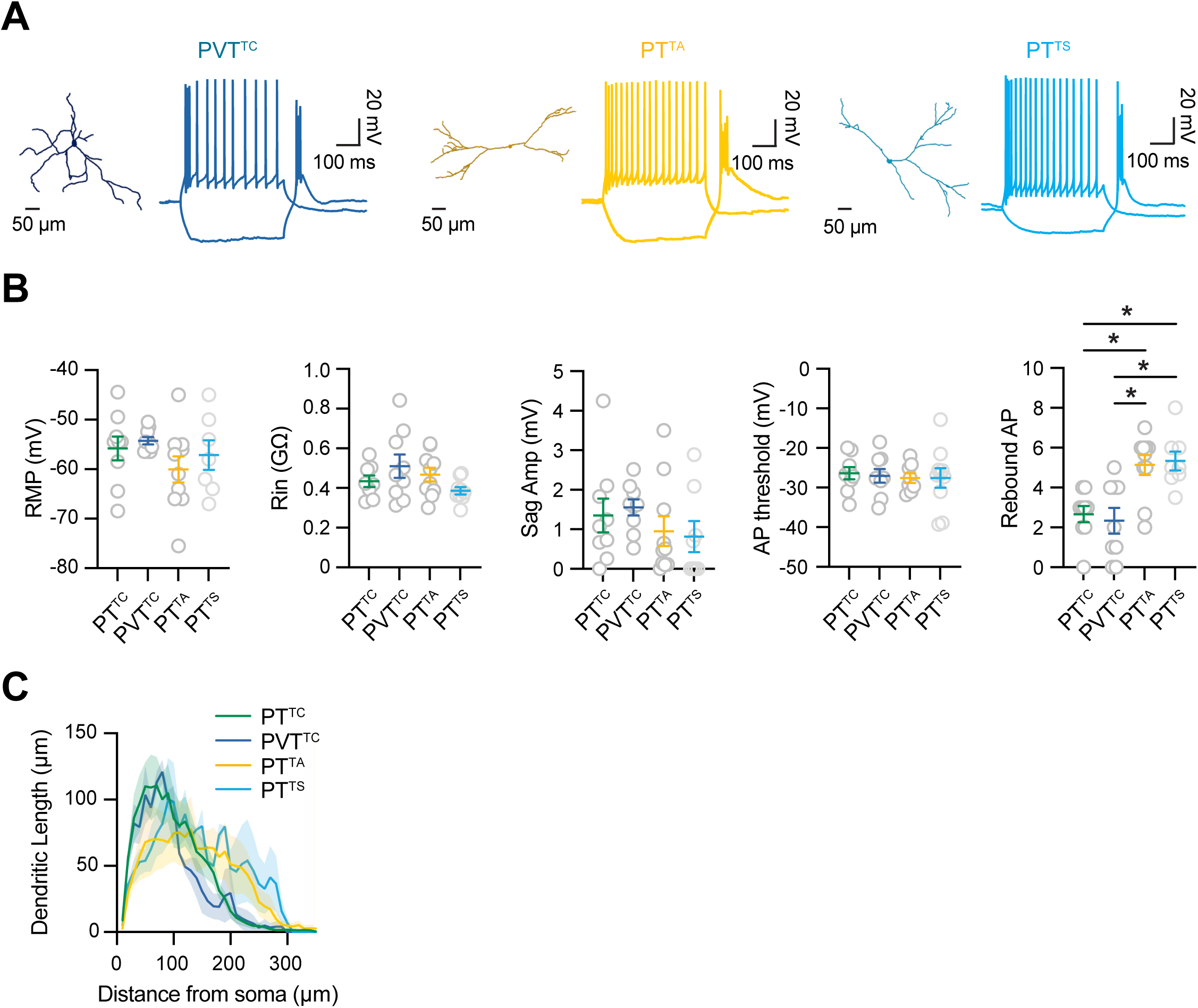
Additional analysis of intrinsic properties of projection neurons in PT and PVT. **(A)** Representative dendritic reconstructions and membrane voltage responses of TC cells in PVT, TS cells in PT, and TA cells in PT. Scale bars = 50 µm and 20 mV x 100 ms. **(B)** Summary of resting membrane potential, input resistance, sag amplitude, AP threshold, and rebound firing of TC cells in PT, TC cells in PVT, TS cells in PT, and TA cells in PT (n = 7-9 cells per group, N = 5-6 mice). **(C)** Summary of Sholl analysis of dendrites for the 4 different cell types (n = 5-9 cells per group, N = 3-5 mice). Values are mean ± SEM. * = *p* < 0.05 *(Related to Figure 2)*

**Figure S3:**
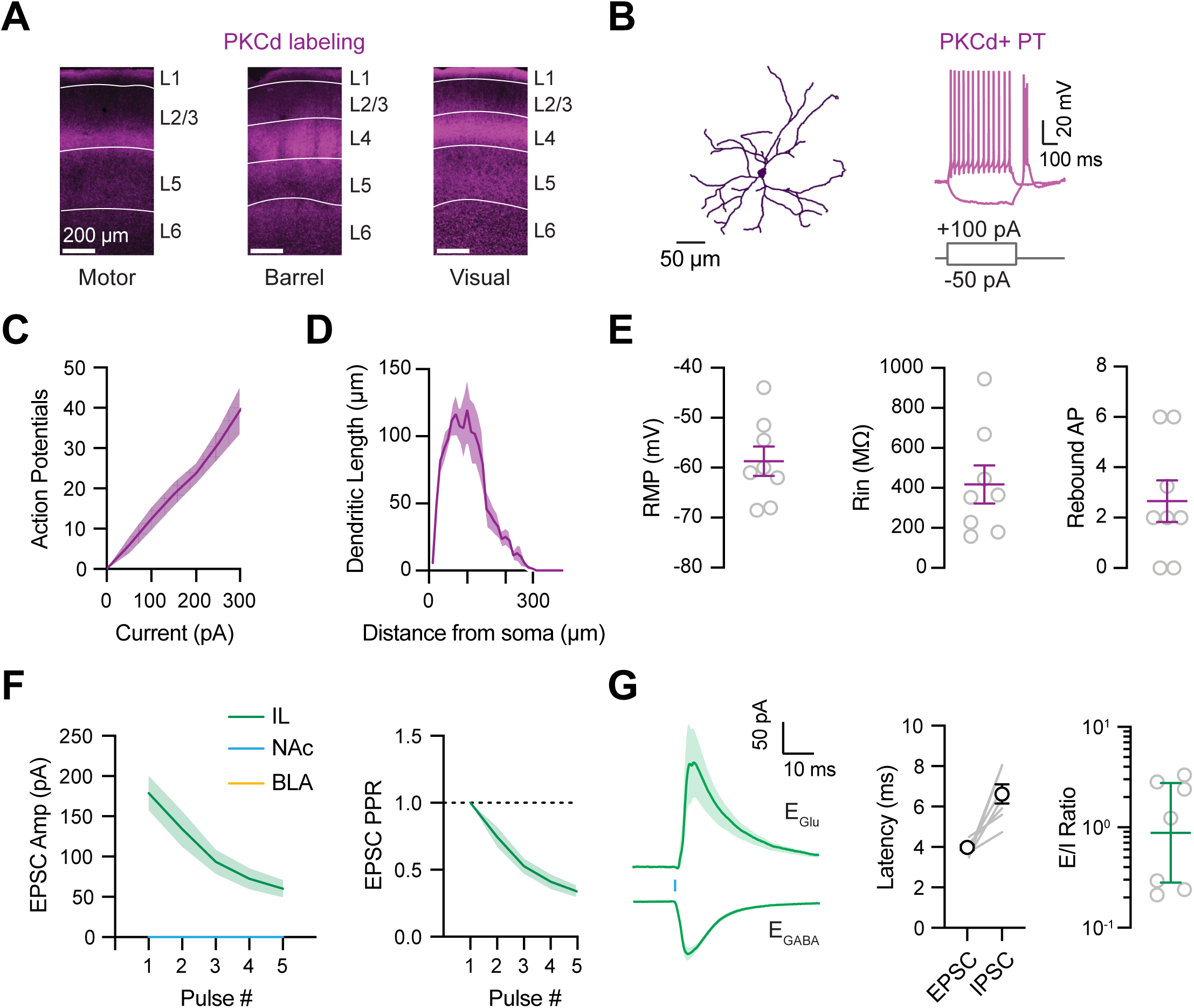
Additional properties of PKCd+ cells and their connections in mPFC. (A) Confocal images of PKCd IHC labeling in motor cortex (left), barrel cortex (middle), and visual cortex (right). Scale bars = 200 µm. (B) Representative dendritic reconstructions and membrane voltage responses of PKCd+ cells in PT. Scale bars = 50 µm and 20 mV x 100 ms. (C) Summary of and frequency-current (F-I) curves (left) (n = 8 cells, N = 3 mice) (D) Summary of Sholl analysis of dendritic length of PKCd+ cells (n = 6 cells, N = 3 mice). (E) Summary of resting membrane potential (RMP), input resistance (Rin), and rebound AP, of PKCd+ cells in PT (n = 8 cells, N = 3 mice). (F) *Left,* Summary of PKCd+ PT-evoked EPSCs in IL, NAc, and BLA as a function of pulse number (n = 8-9 cells per area, N = 5 mice). *Right,* PPR of PKCd+ PT-evoked EPSCs in IL. (G) *Left*, Average PKCd+ PT-evoked EPSCs recorded at E_GABA_ and IPSCs recorded at E_glu_ at IL L2/3 pyramidal cells. Scale bars = 50 pA x 10 ms. *Middle*, Summary of latency from stimulation to EPSC and IPSC, indicating feed-forward inhibition. *Right*, Summary of excitation / inhibition (E/I) ratio (n = 7 cells, N = 5 mice). Values are presented as mean ± SEM, or geometric means ± 95% CI (panel E). *(Related to Figure 3)*

**Figure S4:**
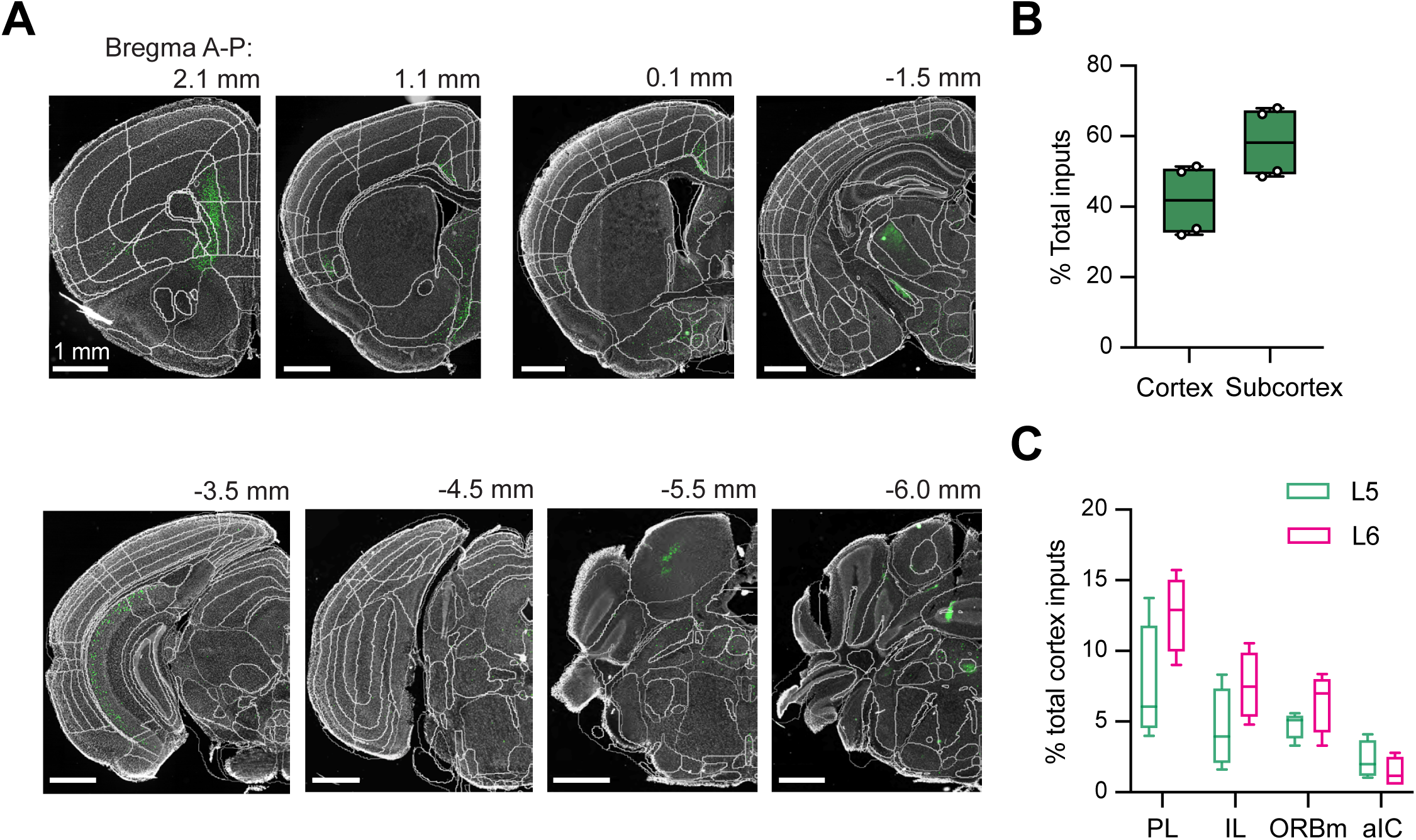
Details of presynaptic inputs to PKCd+ cells in the paratenial thalamus. **(A)** Representative images of RV-labeled presynaptic input cells (green) across the A-P axis of the brain, with distances reported relative to Bregma. Scale bars = 1 mm. **(B)** Summary of percent of total cortical vs. subcortical input cells (N = 4 mice). **(C)** Summary of percent of total cortical input cells residing in L5 and L6 (N = 4 mice). Box and whisker plots represent median and minimum to maximum. (*Related to Figure 4)*

**Figure S5:**
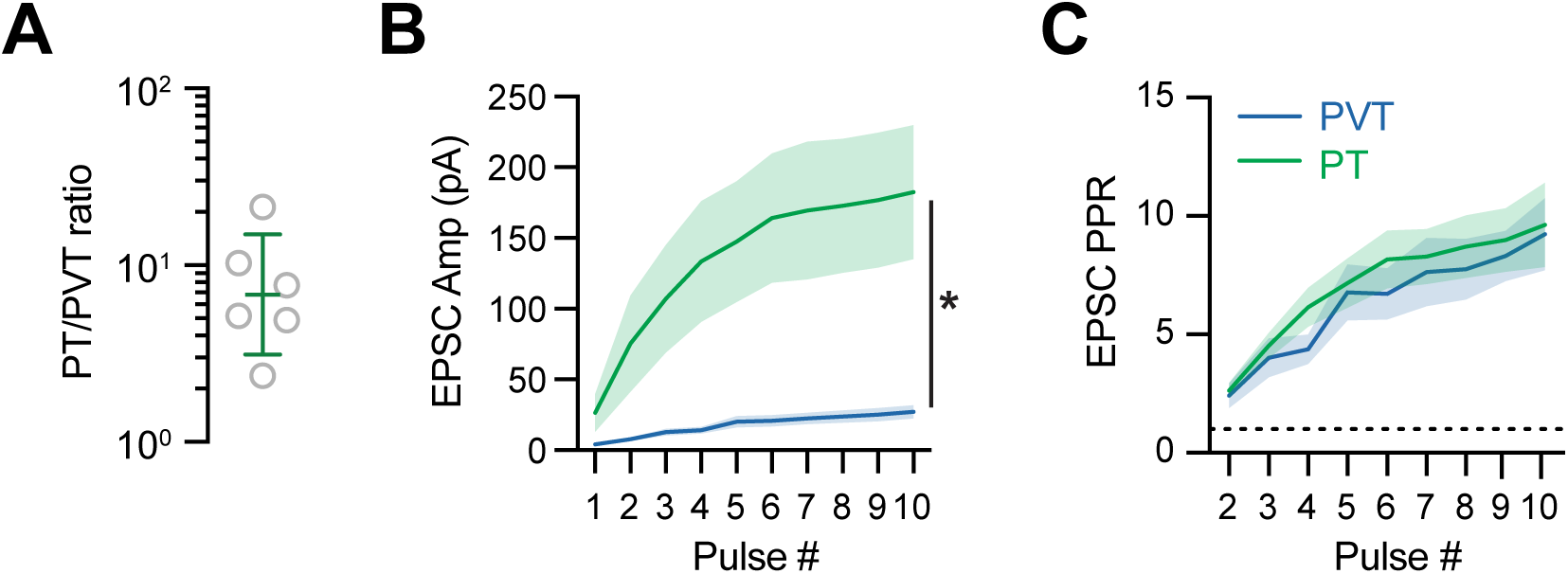
Further quantification if mPFC-evoked EPSCs at TC cells in PT and PVT. **(A)** Summary of PT/PVT ratio of mPFC-evoked EPSCs at TC cells (n = 6 pairs, N = 4 mice). **(B)** Summary of mPFC-evoked EPSC amplitudes at pairs of TC cells in PT and PVT as a function of stimulation pulse number. **(C)** Summary of paired-pulse ratio (PPR) of EPSCs. Values are presented as mean ± SEM. (*Related to Figure 5)*

**Figure S6:**
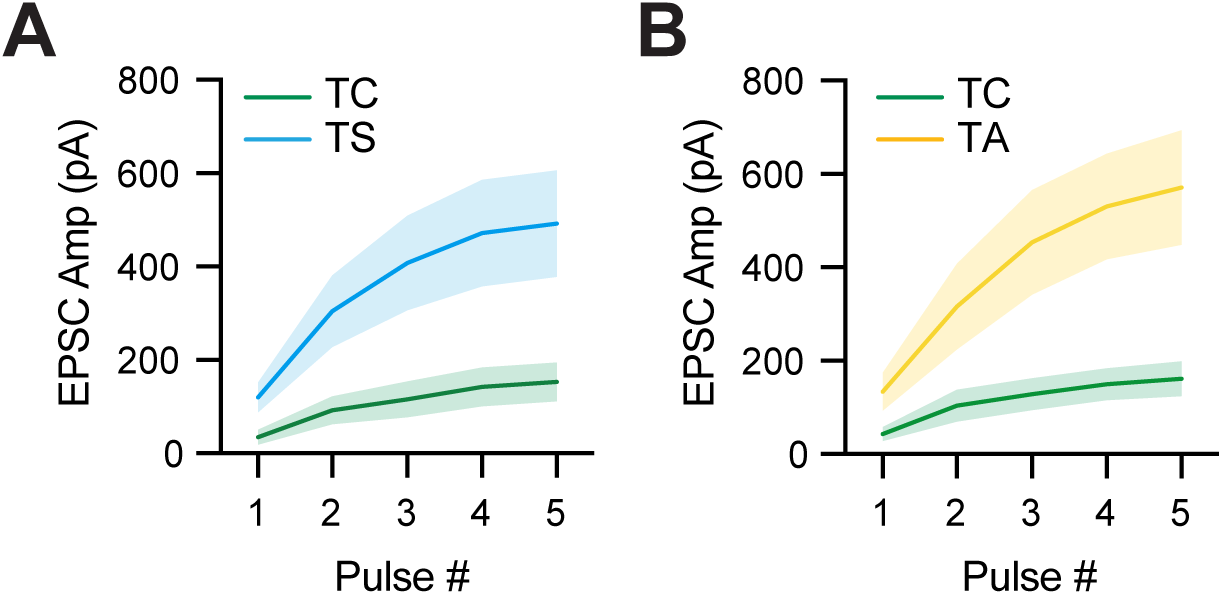
Further quantification if mPFC-evoked EPSCs at TC, TS, and TA cells. **(A)** Summary of mPFC-evoked EPSC amplitudes at pairs of TC and TS cells as a function of stimulation pulse number. **(B)** Similar to (A) for pairs of TC and TA cells. Values are presented as mean ± SEM. (*Related to Figure 6*)

**Figure S7:**
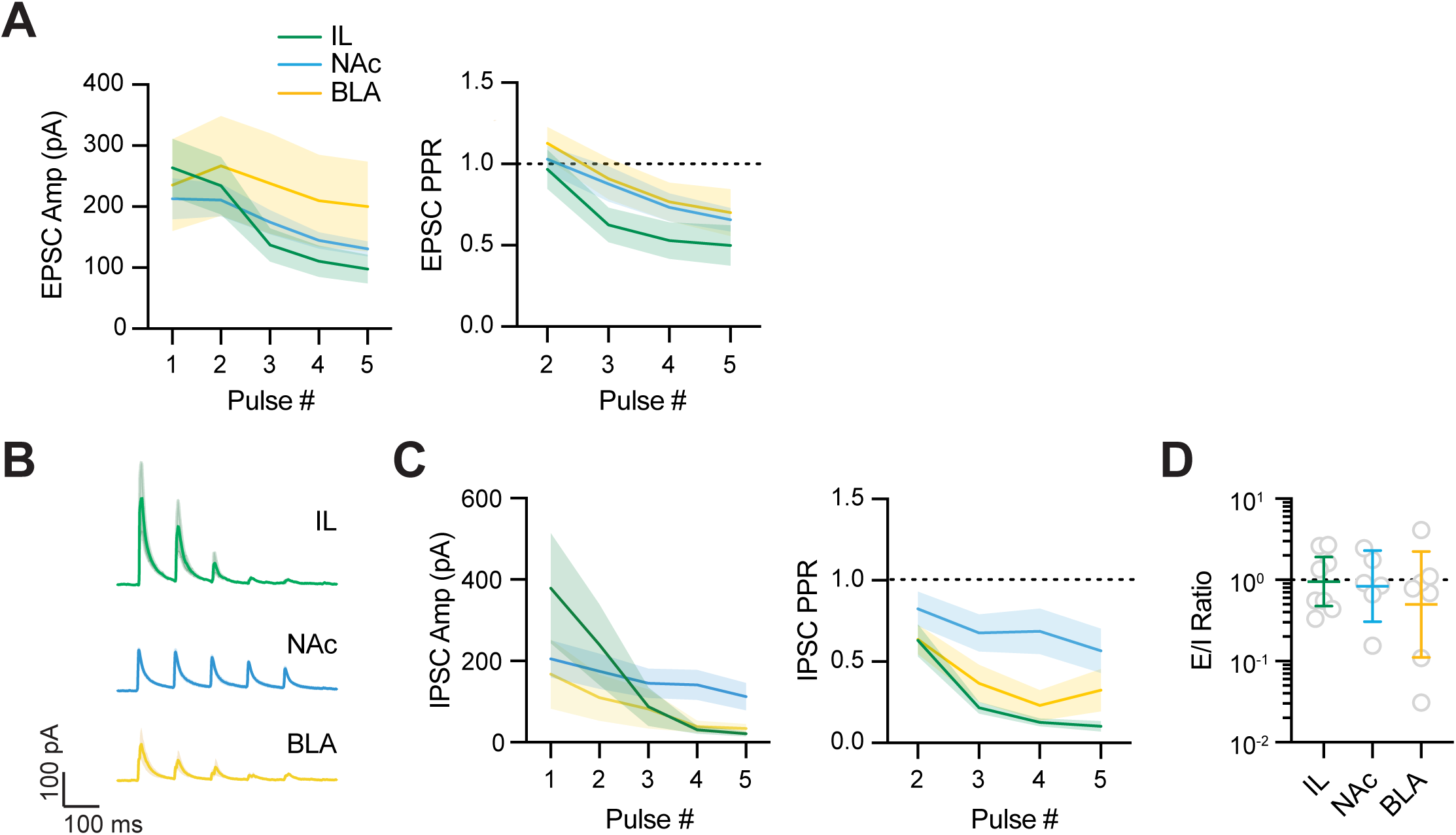
PT-evoked excitation and inhibition in cortical and subcortical targets. (A) *Left,* Summary of PT-evoked EPSC amplitudes in IL, NAc, and BLA as a function of stimulation pulse number (n = 7-8 cells per area, N = 5 mice). *Right,* Summary of PPR of EPSCs. (B) Average PT-evoked IPSCs recorded at E_glu_ in IL L2/3 pyramidal cells (top), NAc MSNs (middle), and BLA pyramidal cells (bottom). Scale bars = 100 pA x 100 ms. (C) *Left,* Summary of PT-evoked IPSC amplitudes in IL, NAc, and BLA as a function of stimulation pulse number (n = 5-6 cells per group). *Right,* Summary of PPR of IPSCs. (D) Summary of excitation / inhibition (E/I) ratio in IL, NAc, and BLA. Values are presented as mean ± SEM or geometric means ± 95% CI. (*Related to Figure 7*)

## REFERENCES

Anastasiades, P.G., Boada, C. & Carter, A.G. (2019) Cell-Type-Specific D1 Dopamine Receptor Modulation of Projection Neurons and Interneurons in the Prefrontal Cortex. Cereb Cortex, 29, 3224–3242.

Anastasiades, P.G. & Carter, A.G. (2021) Circuit organization of the rodent medial prefrontal cortex. Trends Neurosci, 44, 550–563.

Anastasiades, P.G., Collins, D.P. & Carter, A.G. (2021) Mediodorsal and Ventromedial Thalamus Engage Distinct L1 Circuits in the Prefrontal Cortex. Neuron, 109, 314–330 e314.

Andolina, I.M., Jones, H.E., Wang, W. & Sillito, A.M. (2007) Corticothalamic feedback enhances stimulus responseprecision in the visual system. Proc. Natl. Acad. Sci. U.S.A., 104, 1685–1690.

Aquino-Miranda, G., Jalloul, D., Zhang, X.O., Li, S., Kirouac, G.J., Beierlein, M. & Do Monte, F.H. (2024) Functional properties of corticothalamic circuits targeting paraventricular thalamic neurons. Neuron, 112, 4060–4080 e4067.

Bartlett, E.L. & Smith, P.H. (2002) Effects of paired-pulse and repetitive stimulation on neurons in the rat medial geniculate body. Neuroscience, 113, 957–974.

Berendse, H.W. & Groenewegen, H.J. (1991) Restricted cortical termination fields of the midline and intralaminar thalamic nuclei in the rat. Neuroscience, 42, 73–102.

Biro, L., Buday, Z., Kota, K., Lorincz, S. & Acsady, L. (2025) Convergence and Segregation of Excitatory and Inhibitory Afferents in the Paraventricular Thalamic Nucleus. J Neurosci, 45.

Born, G., Schneider-Soupiadis, F.A., Erisken, S., Vaiceliunaite, A., Lao, C.L., Mobarhan, M.H., Spacek, M.A., Einevoll, G.T. & Busse, L. (2021) Corticothalamic feedback sculpts visual spatial integration in mouse thalamus. Nat Neurosci, 24, 1711–1720.

Callaway, E.M. & Luo, L. (2015) Monosynaptic Circuit Tracing with Glycoprotein-Deleted Rabies Viruses. J Neurosci, 35, 8979–8985.

Castro-Alamancos, M.A. & Connors, B.W. (1997) Thalamocortical synapses. Prog. Neurobiol., 51, 581–606.

Chen, L., Nagaraja, C., Daniels, S., Fisk, Z.A., Dvorak, R., Meyerdirk, L., Steiner, J.A., Escobar Galvis, M.L., Henderson, M.X., Rousseaux, M.W.C., Brundin, P. & Chu, H.Y. (2022) Synaptic location is a determinant of the detrimental effects of alpha-synuclein pathology to glutamatergic transmission in the basolateral amygdala. Elife, 11.

Chen, S.S., H-S (1990) Afferent connections of the thalamic paraventricular and parataenial nuclei in the rat — a retrograde tracing study with iontophoretic application of Fluoro-Gold. Brain Research, 522, 1–6.

Clark, A.M., Leroy, F., Martyniuk, K.M., Feng, W., McManus, E., Bailey, M.R., Javitch, J.A., Balsam, P.D. & Kellendonk, C. (2017) Dopamine D2 Receptors in the Paraventricular Thalamus Attenuate Cocaine Locomotor Sensitization. eNeuro, 4.

Collins, D.P., Anastasiades, P.G., Marlin, J.J. & Carter, A.G. (2018) Reciprocal Circuits Linking the Prefrontal Cortex with Dorsal and Ventral Thalamic Nuclei. Neuron, 98, 366–379 e364.

Crandall, S.R., Cruikshank, S.J. & Connors, B.W. (2015) A corticothalamic switch: controlling the thalamus with dynamic synapses. Neuron, 86, 768–782.

Cruikshank, S.J., Ahmed, O.J., Stevens, T.R., Patrick, S.L., Gonzalez, A.N., Elmaleh, M. & Connors, B.W. (2012) Thalamic control of layer 1 circuits in prefrontal cortex. J Neurosci, 32, 17813–17823.

Cruikshank, S.J., Urabe, H., Nurmikko, A.V. & Connors, B.W. (2010) Pathway-specific feedforward circuits between thalamus and neocortex revealed by selective optical stimulation of axons. Neuron, 65, 230–245.

de Jong, A.P. & Verhage, M. (2009) Presynaptic signal transduction pathways that modulate synaptic transmission. Curr Opin Neurobiol, 19, 245–253.

Ding, J., Peterson, J.D. & Surmeier, D.J. (2008) Corticostriatal and thalamostriatal synapses have distinctive properties. J Neurosci, 28, 6483–6492.

Do-Monte, F.H., Quinones-Laracuente, K. & Quirk, G.J. (2015) A temporal shift in the circuits mediating retrieval of fear memory. Nature, 519, 460–463.

Dolleman-van der Weel, M.J., Griffin, A.L., Ito, H.T., Shapiro, M.L., Witter, M.P., Vertes, R.P. & Allen, T.A. (2019) The nucleus reuniens of the thalamus sits at the nexus of a hippocampus and medial prefrontal cortex circuit enabling memory and behavior. Learn Mem, 26, 191–205.

Dong, X., Li, S. & Kirouac, G.J. (2017) Collateralization of projections from the paraventricular nucleus of the thalamus to the nucleus accumbens, bed nucleus of the stria terminalis, and central nucleus of the amygdala. Brain Struct Funct, 222, 3927–3943.

Finch, D.M. (1996) Neurophysiology of converging synaptic inputs from the rat prefrontal cortex, amygdala, midline thalamus, and hippocampal formation onto single neurons of the caudate/putamen and nucleus accumbens. Hippocampus, 6, 495–512.

Fujiyama, F., Unzai, T., Nakamura, K., Nomura, S. & Kaneko, T. (2006) Difference in organization of corticostriatal and thalamostriatal synapses between patch and matrix compartments of rat neostriatum. Eur J Neurosci, 24, 2813–2824.

Gabbott, P.L., Warner, T.A., Jays, P.R., Salway, P. & Busby, S.J. (2005) Prefrontal cortex in the rat: projections to subcortical autonomic, motor, and limbic centers. J Comp Neurol, 492, 145–177.

Gao, C., Gohel, C.A., Leng, Y., Ma, J., Goldman, D., Levine, A.J. & Penzo, M.A. (2023) Molecular and spatial profiling of the paraventricular nucleus of the thalamus. Elife, 12.

Gao, C., Leng, Y., Ma, J., Rooke, V., Rodriguez-Gonzalez, S., Ramakrishnan, C., Deisseroth, K. & Penzo, M.A. (2020) Two genetically, anatomically and functionally distinct cell types segregate across anteroposterior axis of paraventricular thalamus. Nat Neurosci, 23, 217–228.

Gimenez-Amaya, J.M., McFarland, N.R., de las Heras, S. & Haber, S.N. (1995) Organization of thalamic projections to the ventral striatum in the primate. J Comp Neurol, 354, 127–149.

Groenewegen, H.J. & Berendse, H.W. (1994) The specificity of the ‘nonspecific’midline and intralaminar thalamic nuclei. Trends in Neurosciences, 17, 52–57.

Halassa, M.M. & Sherman, S.M. (2019) Thalamocortical Circuit Motifs: A General Framework. Neuron, 103, 762–770.

Harris, R.M. (1986) Morphology of physiologically identified thalamocortical relay neurons in the rat ventrobasal thalamus. J Comp Neurol, 251, 491–505.

Haubensak, W., Kunwar, P.S., Cai, H., Ciocchi, S., Wall, N.R., Ponnusamy, R., Biag, J., Dong, H.W., Deisseroth, K., Callaway, E.M., Fanselow, M.S., Luthi, A. & Anderson, D.J. (2010) Genetic dissection of an amygdala microcircuit that gates conditioned fear. Nature, 468, 270–276.

Hsu, D.T. & Price, J.L. (2007) Midline and intralaminar thalamic connections with the orbital and medial prefrontal networks in macaque monkeys. J Comp Neurol, 504, 89–111.

Hua, R., Wang, X., Chen, X., Wang, X., Huang, P., Li, P., Mei, W. & Li, H. (2018) Calretinin Neurons in the Midline Thalamus Modulate Starvation-Induced Arousal. Curr Biol, 28, 3948–3959 e3944.

Ibrahim, B.A., Murphy, C.A., Yudintsev, G., Shinagawa, Y., Banks, M.I. & Llano, D.A. (2021) Corticothalamic gating of population auditory thalamocortical transmission in mouse. Elife, 10.

Iglesias, A.G. & Flagel, S.B. (2021) The Paraventricular Thalamus as a Critical Node of Motivated Behavior via the Hypothalamic-Thalamic-Striatal Circuit. Front Integr Neurosci, 15, 706713.

Jones, E.G. (1998) VIEWPOINT: THE CORE AND MATRIX OF THALAMIC ORGANIZATION. Neuroscience, 85, 331–345.

Kamalova, A., Manoocheri, K., Liu, X., Casello, S.M., Huang, M., Baimel, C., Jang, E.V., Anastasiades, P.G., Collins, D.P. & Carter, A.G. (2024) CCK+ Interneurons Contribute to Thalamus-Evoked Feed-Forward Inhibition in the Prelimbic Prefrontal Cortex. J Neurosci, 44.

Kessler, S., Labouebe, G., Croizier, S., Gaspari, S., Tarussio, D. & Thorens, B. (2021) Glucokinase neurons of the paraventricular nucleus of the thalamus sense glucose and decrease food consumption. iScience, 24, 103122.

Kirouac, G.J. (2015) Placing the paraventricular nucleus of the thalamus within the brain circuits that control behavior. Neurosci Biobehav Rev, 56, 315–329.

Kooiker, C.L., Chen, Y., Birnie, M.T. & Baram, T.Z. (2023) Genetic Tagging Uncovers a Robust, Selective Activation of the Thalamic Paraventricular Nucleus by Adverse Experiences Early in Life. Biol Psychiatry Glob Open Sci, 3, 746–755.

Levey, A.I., Hallanger, A.E. & Wainer, B.H. (1987) Choline acetyltransferase immunoreactivity in the rat thalamus. J Comp Neurol, 257, 317–332.

Li, H., Namburi, P., Olson, J.M., Borio, M., Lemieux, M.E., Beyeler, A., Calhoon, G.G., Hitora-Imamura, N., Coley, A.A., Libster, A., Bal, A., Jin, X., Wang, H., Jia, C., Choudhury, S.R., Shi, X., Felix-Ortiz, A.C., de la Fuente, V., Barth, V.P., King, H.O., Izadmehr, E.M., Revanna, J.S., Batra, K., Fischer, K.B., Keyes, L.R., Padilla-Coreano, N., Siciliano, C.A., McCullough, K.M., Wichmann, R., Ressler, K.J., Fiete, I.R., Zhang, F., Li, Y. & Tye, K.M. (2022) Neurotensin orchestrates valence assignment in the amygdala. Nature, 608, 586–592.

Li, S.H., Li, S. & Kirouac, G.J. (2024) Analysis of Monosynaptic Inputs to Thalamic Paraventricular Nucleus Neurons Innervating the Shell of the Nucleus Accumbens and Central Extended Amygdala. Neuroscience, 537, 151–164.

Linley, S.B., Rojas, A.K.P. & Vertes, R.P. (2025) Afferent Projections to the Paratenial Nucleus of the Dorsal Midline Thalamus. J Comp Neurol, 533, e70082.

Liu, X. & Carter, A.G. (2018) Ventral Hippocampal Inputs Preferentially Drive Corticocortical Neurons in the Infralimbic Prefrontal Cortex. J Neurosci, 38, 7351–7363.

Llinás, R. & Jahnsen, H. (1982) Electrophysiology of mammalian thalamic neurones in vitro. Nature, 297, 406–408.

Lui, J.H., Nguyen, N.D., Grutzner, S.M., Darmanis, S., Peixoto, D., Wagner, M.J., Allen, W.E., Kebschull, J.M., Richman, E.B., Ren, J., Newsome, W.T., Quake, S.R. & Luo, L. (2021) Differential encoding in prefrontal cortex projection neuron classes across cognitive tasks. Cell, 184, 489–506 e426.

Ma, J., du Hoffmann, J., Kindel, M., Beas, B.S., Chudasama, Y. & Penzo, M.A. (2021) Divergent projections of the paraventricular nucleus of the thalamus mediate the selection of passive and active defensive behaviors. Nat Neurosci, 24, 1429–1440.

Ma, J., O’Malley, J.J., Kreiker, M., Leng, Y., Khan, I., Kindel, M. & Penzo, M.A. (2024) Convergent direct and indirect cortical streams shape avoidance decisions in mice via the midline thalamus. Nat Commun, 15, 6598.

Matyas, F., Komlosi, G., Babiczky, A., Kocsis, K., Bartho, P., Barsy, B., David, C., Kanti, V., Porrero, C., Magyar, A., Szucs, I., Clasca, F. & Acsady, L. (2018) A highly collateralized thalamic cell type with arousal-predicting activity serves as a key hub for graded state transitions in the forebrain. Nat Neurosci, 21, 1551–1562.

McAllister, J.P. & Wells, J. (1981) The structural organization of the ventral posterolateral nucleus in the rat. J Comp Neurol, 197, 271–301.

McGinty, J.F. & Otis, J.M. (2020) Heterogeneity in the Paraventricular Thalamus: The Traffic Light of Motivated Behaviors. Front Behav Neurosci, 14, 590528.

Oberlaender, M., de Kock, C.P., Bruno, R.M., Ramirez, A., Meyer, H.S., Dercksen, V.J., Helmstaedter, M. & Sakmann, B. (2012) Cell type-specific three-dimensional structure of thalamocortical circuits in a column of rat vibrissal cortex. Cereb Cortex, 22, 2375–2391.

Olsen, S.R., Bortone, D.S., Adesnik, H. & Scanziani, M. (2012) Gain control by layer six in cortical circuits of vision. Nature, 483, 47–52.

Otis, J.M., Namboodiri, V.M., Matan, A.M., Voets, E.S., Mohorn, E.P., Kosyk, O., McHenry, J.A., Robinson, J.E., Resendez, S.L., Rossi, M.A. & Stuber, G.D. (2017) Prefrontal cortex output circuits guide reward seeking through divergent cue encoding. Nature, 543, 103–107.

Paniccia, J.E., Vollmer, K.M., Green, L.M., Grant, R.I., Winston, K.T., Buchmaier, S., Westphal, A.M., Clarke, R.E., Doncheck, E.M., Bordieanu, B., Manusky, L.M., Martino, M.R., Ward, A.L., Rinker, J.A., McGinty, J.F., Scofield, M.D. & Otis, J.M. (2024) Restoration of a paraventricular thalamo-accumbal behavioral suppression circuit prevents reinstatement of heroin seeking. Neuron, 112, 772–785 e779.

Penzo, M.A., Robert, V., Tucciarone, J., De Bundel, D., Wang, M., Van Aelst, L., Darvas, M., Parada, L.F., Palmiter, R.D., He, M., Huang, Z.J. & Li, B. (2015) The paraventricular thalamus controls a central amygdala fear circuit. Nature, 519, 455–459.

Petreanu, L., Mao, T., Sternson, S.M. & Svoboda, K. (2009) The subcellular organization of neocortical excitatory connections. Nature, 457, 1142–1145.

Reichova, I. & Sherman, S.M. (2004) Somatosensory corticothalamic projections: distinguishing drivers from modulators. J Neurophysiol, 92, 2185–2197.

Room, P., Russchen, F.T., Groenewegen, H.J. & Lohman, A.H. (1985) Efferent connections of the prelimbic (area 32) and the infralimbic (area 25) cortices: an anterograde tracing study in the cat. J Comp Neurol, 242, 40–55.

Schaeuble, D. & Myers, B. (2022) Cortical-Hypothalamic Integration of Autonomic and Endocrine Stress Responses. Front Physiol, 13, 820398.

Schmitt, L.I., Wimmer, R.D., Nakajima, M., Happ, M., Mofakham, S. & Halassa, M.M. (2017) Thalamic amplification of cortical connectivity sustains attentional control. Nature, 545, 219–223.

Shima, Y., Skibbe, H., Sasagawa, Y., Fujimori, N., Iwayama, Y., Isomura-Matoba, A., Yano, M., Ichikawa, T., Nikaido, I., Hattori, N. & Kato, T. (2023) Distinctiveness and continuity in transcriptome and connectivity in the anterior-posterior axis of the paraventricular nucleus of the thalamus. Cell Rep, 42, 113309.

Smeal, R.M., Keefe, K.A. & Wilcox, K.S. (2008) Differences in excitatory transmission between thalamic and cortical afferents to single spiny efferent neurons of rat dorsal striatum. Eur J Neurosci, 28, 2041–2052.

Szinyei, C., Heinbockel, T., Montagne, J. & Pape, H.C. (2000) Putative Cortical and Thalamic Inputs Elicit Convergent Excitation in a Population of GABAergic Interneurons of the Lateral Amygdala. J Neurosci, 20, 8909–8915.

Tang, Q.Q., Wu, Y., Tao, Q., Shen, Y., An, X., Liu, D. & Xu, Z. (2024) Direct paraventricular thalamus-basolateral amygdala circuit modulates neuropathic pain and emotional anxiety. Neuropsychopharmacology, 49, 455–466.

Theyel, B.B., Llano, D.A. & Sherman, S.M. (2010) The corticothalamocortical circuit drives higher-order cortex in the mouse. Nat Neurosci, 13, 84–88.

Vaasjo, L.O., Han, X., Thurmon, A.N., Tiemroth, A.S., Berndt, H., Korn, M., Figueroa, A., Reyes, R., Feliciano-Ramos, P.A. & Galazo, M.J. (2022) Characterization and manipulation of Corticothalamic neurons in associative cortices using Syt6-Cre transgenic mice. J Comp Neurol, 530, 1020–1048.

Vertes, R.P. (2004) Differential projections of the infralimbic and prelimbic cortex in the rat. Synapse, 51, 32–58.

Vertes, R.P. & Hoover, W.B. (2008) Projections of the paraventricular and paratenial nuclei of the dorsal midline thalamus in the rat. J Comp Neurol, 508, 212–237.

Vertes, R.P., Linley, S.B. & Hoover, W.B. (2015) Limbic circuitry of the midline thalamus. Neurosci Biobehav Rev, 54, 89–107.

Viena, T.D., Rasch, G.E., Silva, D. & Allen, T.A. (2021) Calretinin and calbindin architecture of the midline thalamus associated with prefrontal-hippocampal circuitry. Hippocampus, 31, 770–789.

Wang, Q., Ding, S.L., Li, Y., Royall, J., Feng, D., Lesnar, P., Graddis, N., Naeemi, M., Facer, B., Ho, A., Dolbeare, T., Blanchard, B., Dee, N., Wakeman, W., Hirokawa, K.E., Szafer, A., Sunkin, S.M., Oh, S.W., Bernard, A., Phillips, J.W., Hawrylycz, M., Koch, C., Zeng, H., Harris, J.A. & Ng, L. (2020) The Allen Mouse Brain Common Coordinate Framework: A 3D Reference Atlas. Cell, 181, 936–953 e920.

Yamamuro, K., Bicks, L.K., Leventhal, M.B., Kato, D., Im, S., Flanigan, M.E., Garkun, Y., Norman, K.J., Caro, K., Sadahiro, M., Kullander, K., Akbarian, S., Russo, S.J. & Morishita, H. (2020) A prefrontal-paraventricular thalamus circuit requires juvenile social experience to regulate adult sociability in mice. Nat Neurosci, 23, 1240–1252.

Yao, Z., van Velthoven, C.T.J., Nguyen, T.N., Goldy, J., Sedeno-Cortes, A.E., Baftizadeh, F., Bertagnolli, D., Casper, T., Chiang, M., Crichton, K., Ding, S.L., Fong, O., Garren, E., Glandon, A., Gouwens, N.W., Gray, J., Graybuck, L.T., Hawrylycz, M.J., Hirschstein, D., Kroll, M., Lathia, K., Lee, C., Levi, B., McMillen, D., Mok, S., Pham, T., Ren, Q., Rimorin, C., Shapovalova, N., Sulc, J., Sunkin, S.M., Tieu, M., Torkelson, A., Tung, H., Ward, K., Dee, N., Smith, K.A., Tasic, B. & Zeng, H. (2021) A taxonomy of transcriptomic cell types across the isocortex and hippocampal formation. Cell, 184, 3222–3241 e3226.

Zhu, Y., Wienecke, C.F., Nachtrab, G. & Chen, X. (2016) A thalamic input to the nucleus accumbens mediates opiate dependence. Nature, 530, 219–222.

Zingg, B., Chou, X.L., Zhang, Z.G., Mesik, L., Liang, F., Tao, H.W. & Zhang, L.I. (2017) AAV-Mediated Anterograde Transsynaptic Tagging: Mapping Corticocollicular Input-Defined Neural Pathways for Defense Behaviors. Neuron, 93, 33–47.

